# Functional Impact of Nth-like DNA glycosylase on Mitochondrial Dynamics

**DOI:** 10.1101/2024.10.10.617535

**Authors:** Lisa Hubers, Alexander Myhr Skjetne, Luisa Luna, Yohan Lefol, Solveig Osnes Lund, Ane Marit Wågbø, Xavier Renaudin, Annikka Polster, Anders Knoph Berg-Eriksen, Francisco Jose Naranjo-Galindo, Anna Campalans, Torkild Visnes, Hilde Loge Nilsen, Nicola P. Montaldo

## Abstract

NTH-like DNA glycosylase (NTHL1), a key enzyme in the base excision repair (BER) pathway, has long been considered essential for maintaining both nuclear and mitochondrial genome integrity. In this study, we integrate *in vitro* biochemical assays, *in-cellulo* molecular biology experiments, and bioinformatic analyses to investigate the impact of NTHL1 loss on mitochondrial DNA (mtDNA) stability and function.

Our findings challenge the conventional view by showing that the loss of NTHL1 confers a positive phenotype. Although NTHL1 deletion in human cells results in a significant accumulation of mtDNA lesions, it is unexpectedly accompanied by a substantial increase in mtDNA copy number (CN), which correlates with elevated oxidative phosphorylation (OXPHOS) protein levels and enhanced mitochondrial respiratory capacity. Additionally, NTHL1-knockout cells exhibit larger mitochondria and increased levels of the mitochondrial biogenesis regulator peroxisome proliferator-activated receptor γ coactivator 1α (PGC1α) and the membrane fusion protein Dynamin-like GTPase OPA1. These changes suggest an adaptive response to stress or altered cellular demands, aimed at enhancing mitochondrial function and capacity. Consequently, *NTHL1*-knockout cells showed resistance to 1-methyl-4-phenylpyridinium (MPP+)-induced mitochondrial stress along with increased phosphorylation of eIF2α. Together, this suggests that the loss of NTHL1 may enhance cellular resilience to oxidative stress by activating protective integrated stress response and mitohormesis pathways thereby contributing to cellular survival under adverse conditions. Collectively, our findings position NTHL1 as a key player in mtDNA stability and mitochondrial function, linking its DNA repair activity to the integrated stress response. Its loss triggers a shift from canonical oxidative phosphorylation to stress-adaptive pathways, ultimately enhancing cellular resistance against oxidative stress through the activation of the mitochondrial hormetic response.

## INTRODUCTION

Mitochondria are key organelles that support cellular energy production, metabolism, and signaling. Mitochondrial DNA (mtDNA) encodes 37 essential genes necessary for mitochondrial function and cellular signaling, including critical components of the oxidative phosphorylation (OXPHOS) system, indispensable for cellular respiration and aerobic metabolism^1–3^. Mutations or defects in mtDNA, along with alterations in nuclear genes that regulate its maintenance, are linked to mitochondrial disorders in both children and adults ^4,5^. mtDNA is particularly vulnerable to damage due to its close proximity to the electron transport chain (ETC) and the lack of protective histones, making it more exposed to reactive oxygen species (ROS) generated during cellular respiration. Its integrity is preserved through multiple mechanisms, including DNA repair, mtDNA turnover, copy number redundancy, as well as mitochondrial quality control processes such as fission, fusion, and mitophagy^3,6^.

DNA base lesions occur frequently in cells due to exposure to both endogenous and exogenous DNA-damaging agents^7^. Among the various forms of damage, oxidized bases are particularly detrimental, necessitating efficient repair mechanisms to maintain genomic integrity. Base excision repair (BER) is the primary pathway for repair of such DNA lesions^8^ and serves as predominant repair system in mitochondria, especially given the limited presence of other DNA repair mechanisms within this organelle^3^. DNA glycosylases are a family of enzymes that initiate the BER pathway by recognizing and subsequently excising damaged bases. Bifunctional DNA glycosylases also have the ability to incise the DNA backbone to generate a single-stranded break that is subsequently processed by apurinic/apyrimidinic (AP) endonuclease 1 (APE1). Finally, DNA polymerase β catalyzes the insertion of the missing nucleotides, while DNA ligase III/XRCC1 seals the nick. Various DNA glycosylases have been identified to target specific types of DNA damage, such as 8-oxoguanine, recognized by 8-Oxoguanine DNA Glycosylase 1 (OGG1) and uracil, recognized by Uracil-DNA Glycosylase (UNG).

The human bifunctional NTH-like DNA glycosylase NTHL1 (also referred to as hNTHL1 or NTH1), is the evolutionary conserved homolog of bacterial Endonuclease III (Nth), and excises oxidized pyrimidines like 5-hydroxyuracil (5-ohU) and thymine glycol (Tg)^9^. Recent research suggests a complex interplay between BER, oxidative stress, and mitochondrial dysfunction in neurodegeneration and cancer. Specifically, BER has been implicated in the age-dependent accumulation of single-stranded DNA breaks, contributing to Parkinson’s disease (PD) and Alzheimer’s disease (AD) pathology in *Caenorhabditis elegans (C. elegans)*^10,11^. In these disease models, the *C. elegans* ortholog NTH-1 generates strand breaks during physiological aging and its loss is associated with a protective phenotype characterized by reduced proteotoxicity and activation of cellular defenses that improve overall health as well as cognitive function^10,11^. Similarly, NTHL1 overexpression causes genomic instability and early cellular hallmarks of cancer^12^. At the same time biallelic inactivating variants in the *NTHL1* gene causes NTHL1 tumor syndrome characterized by increased lifetime risk for colorectal cancer^13–16^. Despite these observations, mechanistic insight on the importance of NTHL1 in the context of mammalian aging, neurodegeneration, and cancer remains limited. In particular, the contribution of the mitochondrial NTHL1 function for cellular health is poorly described.

mtDNA copy number (mtDNA-CN) is typically associated with energy production and mitochondrial membrane potential^17^. mtDNA-CN decline during aging, with lower levels being associated with cognitive and physical decline, as well as higher mortality risk^18–20^. Reductions in mtDNA-CN have been implicated in the progression of PD^21,22^ and could serve as a valuable prognostic marker for disease advancement. mtDNA-CN alterations have also been observed across various cancers^23^, with both increases and decreases reported in primary human cancers, potentially contributing to cancer pathogenesis and progression^24^. The exact mechanisms underlying these alterations remain poorly understood particularly the role of DNA repair pathways and specific DNA lesions in regulating mtDNA-CN. Further investigation is needed to clarify these relationships and their impact on neurodegeneration and cancer development.

In this study, we integrate *in vitro* biochemical assays, *in cellulo-*molecular biology experiments, and bioinformatic analyses to unveil the impact of NTHL1 on mtDNA and mitochondrial function. RNA sequencing of CRISPR/Cas9-generated *NTHL1^−/−^* cell lines revealed altered expression of mitochondrial-related genes, particularly those involved in oxidoreductase activity and the ETC, suggesting regulation in key mitochondrial functions such as energy production. This reprogramming was associated with increased mtDNA-CN, elevated mtDNA damage, and reduced TFAM binding to mtDNA, indicating impaired mtDNA packaging. Respirometry assays showed enhanced mitochondrial respiration in *NTHL1^−/−^* cells, supported by increased mitochondrial mass, as confirmed by immunofluorescence and transmission electron microscopy. Notably, *NTHL1^−/−^* cells exhibited greater resistance to mitochondrial stress induced by the complex I inhibitor 1-methyl-4-phenylpyridinium (MPP+), suggesting a mitochondrial-specific stress response. This resilience was linked to activation of the integrated stress response (ISR), characterized by increased phosphorylation of eIF2α, a key regulator of mitochondrial function and stress adaptation.

These findings suggest that NTHL1 loss triggers a compensatory response to mtDNA damage, enhancing mitochondrial function through the activation of adaptive stress response pathways to preserve cellular survival and function under oxidative stress.

## MATERIAL AND METHODS

### Cell lines and cell culture

Human embryonic kidney cells (HEK293, ATCC, CRL-1573) and human osteosarcoma cells (U2OS) were cultured at 37°C under a 5% CO_2_ atmosphere using Dulbecco’s Modified Eagle Medium (DMEM) high glucose (Gibco, 11965092) supplemented with 10% fetal bovine serum (FBS) (Merck, F7524) and 1% penicillin/streptomycin (P/S) (Fischer Scientific, 15140130,).

Near-haploid HAP1 cell lines, derived from the KBM-7 cell line, were cultured at 37°C under the same condition using Iscove’s Modified Dulbecco’s Medium (IMDM) (Gibco, 12440061,) supplemented with 10% FBS and 1% P/S. The HAP1 cell lines included parental clones WT 1.1, WT 1.3 as well as *NTHL1 KO clones 10* and *11* (HZGHC001540c010 and HZGHC001540c011, Horizon Discovery, PerkinElmer).

### Generation of HEK293 Knockout cell lines

Short guide RNA (sgRNA) targeting *NTHL1* gene and potential offtargets were designed using the CRISPOR CRISPR Design tool, and potential off-targets were identified through its predictive algorithm^25^. Alt-R® CRISPR/Cas9 tracrRNA ATTO™550 and Guide (crRNA) were combined in a 1:1 and annealed by heating at 95°C for 5 minutes, followed by a slow cooling process at room temperature (RT). 1 μM of the resulting single guide RNA (sgRNA) was mixed with 1 μM Alt-R® S.p. Cas9 Nuclease V3 (IDT) in Opti-MEM and delivered to HEK293 WT using Lipofectamine™ 3000 Transfection Reagent (Invitrogen™). Upon transfection cells were seeded as single clones into a 96-well plate via serial dilution. Insertion-deletion (Indel) mutations and loss of target protein expression were verified in clonal knockout cell lines through Indel specific PCR, sanger sequencing and immunoblot analysis, respectively. Confirmed HEK293 *NTHL1^-/-^* clones 1, 3, and 15 were here referred as *NTHL1^-/-^* 1, 2, and 3, respectively. Top-exonic off-target sites with a maximum of three base pair differences from the sgRNA were evaluated in the selected clones via Indel-specific PCR and Sanger sequencing.

### PCR, gel electrophoresis and sanger sequencing

Total DNA was extracted using the AllPrep kit (Qiagen, 80204). DNA amplification was performed using GoTaq G2 Master Mix (Promega, M7822) at a 1X concentration, with 0.5 µM forward and reverse primers, in a final volume of 50 µL. The amplification was carried out in a thermal cycler, and the PCR products were resolved on a 1% agarose gel. For Sanger sequencing, the band corresponding to the expected size was visualized under UV light, excised, and purified using the NucleoSpin Gel and PCR Clean-up kit (Macherey-Nagel, 740609.250). The purified PCR product was then sent for sequencing with a single primer located near the region of interest. For indel-specific PCR, the forward primer was designed so that its last base would anneal immediately after the CRISPR-Cas9 cut site. This design ensures that Taq polymerase does not elongate the substrate in most cases of indel mutations, resulting in a significantly reduced or absent amplicon. The absence or presence of an amplicon provides a clear yes/no readout from the gel.

### Mitochondria extraction

Mitochondria were isolated from HEK293 cells using the mitochondria isolation kit from Miltenyi Biotec according to manufacturer’s protocol. Briefly, 1 x 10^7 cells were collected, resuspended in lysis buffer and homogenized using a douce homogenizer. Anti-TOM22 microbeads were used to magnetically label mitochondria, followed by column-based magnetic separation. Mitochondria were used immediately for downstream analysis.

### Transfection

U2OS cells were transfected with the plasmid pcDNA3.1-NTH1-FLAG at final concentration of 100ng/mL using Lipofectamine 2000 (Thermofisher, 11668027) according to manufacturer’s instructions.

### Library Construction, Quality Control, and Sequencing

Total RNA was purified with RNeasy kit (Qiagen, 4368708), and DNaseI digested (Qiagen, 79254) according to the manufacturer’s protocol. Total RNA was used as the input material for RNA sample preparations. RNA library preparation, and sequencing, were carried out by Novogene Europe. Ribosomal RNA (rRNA) was removed from total RNA using specific probes. RNA fragmentation was performed using divalent cations at elevated temperatures in the First Strand Synthesis Reaction Buffer. First-strand cDNA was synthesized using random hexamer primers and reverse transcriptase. The second strand of cDNA was then synthesized by adding buffer and dNTPs, where dTTP was replaced by dUTP. The synthesized double-stranded cDNA was subjected to end repair, A-tailing, adaptor ligation, and PCR enrichment following fragment selection. To select cDNA fragments ranging from 370-420 bp, PCR products were purified using AMPure XP beads, resulting in a strand-specific library. Following library construction, preliminary quantification was performed using Qubit. The size of the inserted fragments was evaluated, and the library’s effective concentration was quantified using reverse transcriptase (RT)-quantitative (q)PCR. The qualified libraries were then pooled and sequenced on Illumina platforms according to the required effective concentration and data output. The original fluorescence image files generated by the Illumina platform were converted into short reads (raw data) through base calling. The resulting short reads were stored in FASTQ format^26^.

### RNA sequencing processing

Quality control was applied to the raw reads to eliminate sequence artifacts, such as adapter contamination, low-quality nucleotides, and unrecognizable nucleotides (N). Quality control was performed using Fastp (v0.23.1)^27^. The quality of clean reads was assessed using FastQC (v0.11.9) prior to alignment. Reads were aligned to the human reference genome (GRCh38) from Ensembl release 109^28^ using STAR (v2.7.10b)^29^, with corresponding gene annotations from the same Ensembl release. Gene-level counts were generated during alignment by enabling the quantMode GeneCounts option in STAR. For downstream analyses, counts from the fourth column of the STAR output, corresponding to the reverse-stranded library orientation, were used.

### RNA sequencing analysis

Counts were normalized and processed using DESeq2 (version 1.40.2)^30^. Genes with an absolute log 2-fold change of 1 or more as well as a false discovery rate smaller than 0.05 were treated as significant differentially expressed genes (DEGs). DEGs were clustered using the PART algorithm ^31^, with parameters of recursion set to 100 and a minimum cluster size of 50 genes. The seed was set to ’123456’ for reproducibility purposes. Each cluster was then submitted to gprofiler (R gprofiler2 version 0.2.2)^32^.

OmnipathR (version 3.12.4) was utilised to find genes associated to ATF4. To mark which genes of the dataset activate the ATF4 transcription factors, genes were filtered using a positive MoR (+1) and a positive log 2 fold change as calculated by the previous differential gene expression analysis.

### Principal Component Analysis (PCA)

PCA was conducted to reduce dimensionality and identify patterns of variation in the expression data. Genes included in the PCA were mitochondrial genes meeting the DE criteria. The analysis was performed using SIMCA 18.0. The PCA scores plot visualizes sample clustering, while the loadings plot identifies key genes contributing to group differences.

### Model Validation

The DModX residual plot was used to evaluate model fit by measuring residual variance. If any samples exceeded the critical limit (Dcrit) this would be flagged as potential outliers. The X/Y overview was used to assess the explanatory (R2X) and predictive (Q2X) power of the PCA model.

### Gene Expression Analysis via RT-qPCR

RNA was purified using the RNeasy kit (Qiagen, 74106) including DNase digestion (Qiagen, 79254) according to manufacturer’s protocol. Subsequently, reverse transcription (RT) of RNA to complementary DNA (cDNA) was carried out using high-capacity cDNA reverse transcription kit (Thermo Fisher, 4368814) according to manufacturer’s instructions. Quantitative PCR (qPCR) was performed in in technical triplicates using Power SYBR PCR Master Mix (Applied Biosystems, 4368708) on a StepOnePlus v2.3 Real-Time PCR System (Applied Biosystems). Relative expression levels were determined using the 2(–ΔΔCt) method, with target gene expression normalized to the housekeeping gene *GAPDH* for nuclear-encoded genes and *B2M* for mitochondrial-encoded genes. Primer sequences used in the qPCR analysis are listed in Supplemental Materials.

### DNA Copy Number Analysis

DNA was isolated from cell extracts using the Qiagen AllPrep kit according to manufacturer’s protocol. To digest any remaining RNA, samples were incubated with 10 μg/mL Ribonuclease A and incubated for 1 hour at 37C. DNA was collected by ethanol precipitation, and qPCR was carried out using Power SYBR PCR Master Mix as above. All primers are listed in Supplemental Materials. To determine the mtDNA/genomic DNA ratio, amplification of individual genes from the mitochondrial genome were compared to a gene from the nuclear genome (*B2M*).

### RADF qPCR DNA Damage Assay

qPCR was performed as described previously, with and without Taq1 restriction enzyme. Primers, which are listed in Supplemental Materials, are specific to the mitochondrial genome and contain a Taq1 restriction site. DNA damage frequency was calculated as (2^-) difference between the Taq1-treated and non-treated genomic DNA.

### Mitochondrial DNA damage via Long-Amplicon PCR

To assess the frequency of DNA lesions capable of inhibiting polymerase activity, we employed the long-amplicon quantitative polymerase chain reaction (LA-QPCR) assay previously described ^33^. Here, the presence of DNA lesions will reduce the efficiency of PCR amplification, with the extent of amplification inversely correlating with the number of lesions within the DNA template. PCR was carried out using LongAmp Hot Start Taq2X Master Mix (New England Biolabs, M0533L). No-template controls (NC) and 50% template controls were included in each experimental run. PCR products were run on a 1% agarose gel and band intensities were quantified using ImageJ software. Each sample’s fluorescence value, corrected for the no-template control, was divided by its respective mtDNA copy number. The so normalized values were averaged and normalized to control sample (WT).

### Whole Cell Extraction

Total proteins were extracted from cell pellets using RIPA lysis buffer containing 10mM Tris-HCl, 150mM NaCl, 0.5mM EDTA, 0.1% SDS, 1% Triton X-100, 1% NaDeoxycholate, 0.1mM PMSF and 1x protease inhibitor cocktail (Roche). Samples were incubated on ice for 30 minutes with frequent pipetting, followed by centrifugation for 10 minutes at 12,000 x g to remove cell debris. Alternatively, cell pellets were resuspended in hypotonic lysis buffer (20 mM Hepes pH 7.9, 2 mM MgCl_2_, 0.2 mM EGTA, 10% glycerol, 2 mM DTT, 0.1 mM PMSF, 1x protease inhibitor cocktail) and incubated 5 minutes on ice. Samples were subjected to three freeze-thaw cycles by snap-freezing in liquid nitrogen followed by thawing in a 37°C water bath for 1 minute. NaCl and NP-40 were added to final concentrations of 140 mM and 0.5%, respectively, and samples were incubated on ice for 20 minutes. Samples were slowly diluted with hypotonic lysis buffer + 140mM NaCl until the final concentration of NP-40 reached 0.2%. Samples were sonicated using a Bioruptor for 5 pulses (30 seconds on, 30 seconds off), and centrifuged for 10 minutes at 12,000 x g. Proteins were quantified using Bradford Assay and used for further analysis.

Total protein extracts used for DNA glycosylase and trapping assays were made from cells lysed in lysis buffer (50 mM MOPS, 200 mM KCl, 0,1% Triton X-100, 1 mM EDTA, 1 mM DTT and 1% Protease Inhibitor Cocktail). After a 30 min incubation on ice, the extracts were frozen in liquid nitrogen/thawed at 30 °C three times. Cell debris was removed by 15 min centrifugation at 13,000 rpm. The protein concentration was measured using Bradford assay for further analysis.

### Immunoblot Analysis

10-40 μg protein extract was combined with 4x Laemmli buffer and incubated for 5 minutes at either 95℃ or 25℃, depending on the antibodies used. Samples were run on a SDS-PAGE gel and transferred to a nitrocellulose membrane. Membranes were blocked in 5% milk followed by incubation with primary antibody, either for 1 hour or overnight. Membranes were incubated with HRP-conjugated secondary antibodies for 30-60 minutes prior to developing. Band intensities were measured using ImageJ and relative protein levels were determined by normalising to Tubulin protein levels. Antibodies used are listed in Supplemental Materials.

### Transmission Electron Microscopy

Briefly, cells were fixed in 2.5% glutaraldehyde, stained in 1% osmium tetroxide, and solidified in 1% agar overnight. Agar containing the cell suspension was subjected to graded dehydration in ethanol and propylene oxide before being embedded in Durcupan resin. Ultrathin sections were cut on a Leica RM UC6 ultramicrotome, contrasted with 1% uranyl acetate and 0.3% lead citrate, and left to dry overnight. Sections were placed on an electron microscopy grid and cellular ultrastructure was visualized on the Tecnai 12 biotwin electron microscope.

### Bioenergetics Analysis

To evaluate cellular bioenergetics, we conducted oxygen consumption rate (OCR) measurements using a Seahorse Xfe96 analyzer (Seahorse Biosciences, MA, USA) via mito stress. Cells were seeded into a Seahorse microplate at a density of 7 × 10^4^ cells/well and subjected to the specified treatments. Following treatment, cells were equilibrated in DMEM lacking sodium bicarbonate, supplemented with either 5 or 35 mM of d-glucose (depending on the treatment), 2 mM of glutamine, and 1 mM of sodium pyruvate. OCR measurements included basal OCR and OCR responses to sequential injections of 1.5 μM oligomycin (an ATP synthase inhibitor targeting mitochondrial complex V), 1 μM FCCP (a mitochondrial uncoupler), and 0.5 μM rotenone/antimycin (inhibitors of mitochondrial complexes I and III, respectively). Parameters such as proton leak, maximal respiration, spare respiratory capacity, non-mitochondrial oxygen consumption, and ATP-linked OCR were analyzed using the mito stress test. OCR values (pmol O₂/min) were normalized to Hoechst 33342 levels.

### DNA glycosylase Activity assay

DNA glycosylase activity was measured using standard assays as previously described^34^. In short, a 30-nt oligonucleotide containing a 5-hydroxyuracil (5-ohU) at position 15 and a 24-nt oligonucleotide containing an uracil (U) at position 14, were ^32^P-labeled at the 5ʹ termini using T4 polynucleotide kinase (NEB, M0201S) and [γ-^32^P] ATP (3000 Ci/mmol) (Revity, NEG502A250UC). They were subsequently annealed with their complementary strand containing a normal base opposite the lesion. Reaction mixtures of 10 μl containing 5-10 fmol of the duplex substrate, reaction buffer (70 mM MOPS, 1 mM DTT, 1 mM EDTA and 5% glycerol) and 10 ug total cell extracts were incubated for 2.5 hours at 37°C. After addition of 10 μl of stop solution (80% formamide, 10 mM EDTA, bromophenol blue and xylene cyanol), the reaction mixtures were incubated for 10 min at 95°C. Samples were separated on 20% denaturing polyacrylamide gels and results analyzed using the Amersham Typhoon and the ImageQuant IQTL 8.2 software (both from Amersham Biosciences).

### NaCNBH₃ mediated trapping of enzyme-substrate intermediates

The Schiff base (SB) covalent intermediate between the bifunctional DNA glycosylase NTHL1 and double-stranded 5-ohU containing oligonucleotide was trapped by its irreversible reduction with sodium borohydride Na[BH_3_(CN)], as previously described^34^. Reaction mixtures containing 5-10 fmol of radiolabelled double-stranded 5-ohU substrate described above were incubated alone or with 10 ug total cell extracts in the presence of 6.4 mM NaCNBH_3_, 35 mM MOPS, 0.5 mM DTT, 0.5 mM EDTA and 2.5% glycerol for 2.5 hours at 37°C in. Reactions were then stopped by adding 4 μl of 4x Laemmli protein sample buffer following by incubation for 10 min at 95°C. Samples were separated on 10% polyacrylamide gels and results analyzed using the Amersham Typhoon IP and the ImageQuant IQTL 8.2 software (both from Amersham Biosciences).

### Chromatin immunoprecipitation (ChIP)

The ChIP assay was performed as previously described^35^ with slight modifications. Briefly, isolated mitochondria were cross-linked in a solution of 1% formaldehyde in PBS for 10 min at RT. The cross-linking reaction was stopped by adding glycine to a final concentration of 0.1 M for 5 min. Mitochondrial pellets were resuspended in SDS lysis buffer (50 mM Tris-HCl pH 8, 10 mM EDTA and 1% SDS) supplemented with protease inhibitors and mtDNA sheared to 200-250 bp DNA fragments by sonication with Bioruptor sonicator (Diagenode) for 12 cycles of 30 s. 20 μg of sheared mtDNA were precleared for 3 h at 4 °C with dynabeads Protein G magnetic beads (10003D, Invitrogen) and then incubated with 6 μl TFAM (ABE483, Millipore-Sigma) or IgG (ab172730, Abcam) antibodies in ChIP buffer (16.7 mM Tris-HCl pH 8, 167 mM NaCl, 1.2 mM EDTA, 0.01% SDS, and 1.1% Triton X-100) overnight at 4 °C. The DNA-protein-antibody complexes were isolated using dynabeads Protein G magnetic beads and washed with low salt wash buffer (16.7 mM Tris-HCl pH 8, 167 mM NaCl, 2 mM EDTA, 0.1% SDS, 1% Triton X), high salt wash buffer (16.7 mM Tris-HCl pH 8, 500 mM NaCl, 2 mM EDTA, 0.1% SDS, 1% Triton X) and LiCl wash buffer (10 mM Tris-HCl pH 8, 250 mM LiCl, 1 mM EDTA, 0.5% NP40, 0.5% Na-deoxycholate) for 10 min each followed by two final washes in TE buffer (10 mM Tris-HCl pH 8, 5 mM EDTA) for 5 min each. Immunoprecipitated DNA was eluted from the beads in 300 μl elution buffer (100 mM, NaHCO_3_, 1% SDS) for 20 min at RT and 10 min at 65 °C. For decrosslinking 15 μl 1 M Tris HCl pH 8, 15 μl 1 M NaCl, 7.5 μl of 0.5 M EDTA, and 1 μl of Proteinase K (20 mg ml^−1^) were added and samples were incubated for 1 hour at 50 °C and 65 °C O/N. Nucleic acids were then purified with phenol/chloroform/isoamyl alcohol, precipitated by centrifugation, washed with 70% ethanol, dried and dissolved in nuclease-free water. Levels of immunoprecipitated DNA is expressed as percentage of input (% input) normalized to WT.

### Cellular proliferation and viability assay

30 µL HEK293 cells were seeded in 384-well plates (1,800 cells/well) and incubated overnight. Cells were subsequently added a dilution series of MPP+ (Abcam, AB144783) dissolved in medium and further incubated for 72 h. Cells were stained with final volumes 5 µg/ml Hoechst 33342 (Thermo Scientific, 62249) and 1 µM Propidium iodide (Invitrogen, P3566) for 30 minutes and imaged. Image acquisition was performed using the ImageXpress Micro Confocal High-Content Imaging System (Molecular Devices) with a 10× CFI Plan Apochromat Lambda objective and single band dichroic filter sets for imaging Hoechst (Ex. 377/54 nm, Em. 447/60 nm) and Propidium iodide (Ex. 560/32 nm, Em. 624/40 nm). Four fields of view were imaged per well (Approx. 72 % well coverage). Image analysis was conducted with the CellProfilerTM Cell Image Analysis Software 4.2.1. Cell numbers were determined as the number of nuclei stained with Hoechst and dead cells were detected as Propidium iodide positive. Cell viability was measured with the CellTiter-Glo® 2.0 Cell Viability Assay (Promega, G9243) at 25 % final concentration using SPARK® Multimode Microplate Reader (Tecan) to measure luminescence. Cell seeding and MPP+ exposure were identical to the cell counting assay described above. For HAP1 cell viability, 3 × 10³ cells per well were seeded into a 96-well plate and incubated for 4 hours. Following this, the cells were treated with MPP+ at concentrations indicated in the figures, or with media alone, for 72 hours. After the treatment, PrestoBlue™ Cell Viability Assay (A13262, Invitrogen) was added to each well, and the plate was incubated for an additional 2 hours. Absorbance was measured at 590 nm. The results presented are averages from two distinct HAP1 clones.

### Immunofluorescence

HEK293 cells were seeded onto glass-bottomed 96-well plates (Greiner Sensoplate, Sigma-Aldrich, M4187-16EA) precoated with 40μg/ml fibronectin (Sigma-Aldrich-Merck, F1141) overnight. For γH2AX staining, cells were treated with 10 µM Camptothecin (CPT, Sigma-Aldrich-Merck, C9911) or 220μM MPP+ for 1 hour at 37°, DMSO served as a control. Following the treatment, cells were pre-extracted with ice-cold 0.2% Triton-X 100 for 2 minutes and fixed with 4% paraformaldehyde for 10 minutes at RT. Subsequently, cells were permeabilized using a methanol:acetone solution (1:1) for 10 minutes at RT and blocked with 1% BSA in PBS for 1 hour at RT. Cells were stained with antibodies for 1 hour, washed (3×5min with PBS), and incubated with secondary antibodies for 1 hour at RT. After secondary antibody incubation, cells were washed (3×5min with PBS) before DNA was stained with 4ʹ,6ʹ-diamidino-2-phenylindole (DAPI) (Sigma-Aldrich-Merck, D9542) at a final concentration of 1μg/ml in PBS for 5 minutes at RT and washed an additional 4 times with PBS. Antibodies used are listed in Supplemental Materials. For MitoView mitochondrial live cell analysis, cells were treated with 100nM MitoView Green dye (Biotium) and 2 drops/mL NucBlue Live Reagent (Invitrogen) for 30 minutes at 37°C and immediately imaged. Images were acquired using ImageXpress Micro Confocal High-Content microscope with a 40x objective and analyzed using ImageJ software.

For immunofluorescence experiments related to figure 1G: U2OS cells were grown on 4 well Ibidi µ-Slides for one day before transfection. 24hr after transfection, cells were incubated in 100nM of MitoTracker Red in warm DMEM for 30 min at 37°C. Then cells were washed with warm DMEM and immediately fixed for 20 min at 37°C with 2% formaldehyde in DMEM, rinsed twice with PBS and permeabilised at RT in PBS 0.1% Triton (X100) for 5 min. Cells were incubated in blocking solution (PBS containing 0.1% Triton-X100, 3% BSA and 1% normal goat serum) at 37°C for 1h. Cells were incubated with primary antibody (anti Flag M2, Sigma-Aldrich F1804) diluted at 1/2000 in blocking solution for 1h at 37°C, washed three times for 5 min in PBS containing 0.1% Triton and further incubated with secondary antibodies (Alexa-488 donkey anti-mouse (Invitrogen A11001) diluted 1/1000 in blocking solution for 1 hr at 37°C. Cells were washed three times for 5 min in PBS containing 0.1% Triton. Nuclear DNA was counterstained with 1 µg/ml DAPI. Image acquisition was performed using a NIKON Ti2 / GATACA, W1 spinning disk microscope with 60X oil immersion objective (Plan Apo N.A. 1.4).

### Statistical analysis

Unless otherwise stated all the experiments presented are representative of n≥3 independent experiments. Data analysis was performed using GraphPad Prism 10.2.2. (GraphPad Software, Inc., La Jolla, CA). Statistical significance was determined by one-way ANOVA with Dunnet’s multiple comparison test (Fig.3A-D, and F; Fig.4B, D, F and H-K; Fig.5D and G; Fig.S1H and K; Fig.S3 A-O; Fig.S4 B-C; Fig.S5A-B) and t-test (Fig S6C). All data represent mean values ± SEM. *P ≤ 0.05; **P ≤ 0.01; ***P ≤ 0.001; ****P ≤ 0.0001; ns - not significant.

## RESULTS

### NTHL1 is active in mitochondria

We employed CRISPR/Cas9 to generate *NTHL1^−/−^* cell lines (Fig.1A), targeting all three *NTHL1* isoforms (Fig.S1A). Following transfection of HEK293 wild type (WT) cells with *NTHL1*-specific sgRNA and Cas9 nuclease, individual clones were isolated and initially screened using an indel-specific PCR (Fig.S1B). Clones lacking amplification in the indel-specific PCR were considered potential knockouts (Fig.S1C) and were further validated by Sanger sequencing to confirm the presence of indel mutations at the target site (Fig.S1D). Immunoblotting verified loss of NTHL1 expression in knockout clones (Fig.1A). To ensure specificity, potential off-target effects were predicted using CRISPOR. Top off-target candidates within exon regions were ruled out via indel-specific PCR and Sanger sequencing of the amplified off-target sites (Fig.S1E-H). Of note our approach ensured the knockout of all three isoforms to avoid any potential compensatory regulation. This was important because, despite isoform 1 being the most highly expressed, the other isoforms were still present and detectable (Fig.S1H-K). To interrogate the consequences of NTHL1 knockout, DNA glycosylase activity and the formation of covalent protein-DNA intermediates were evaluated. A significant reduction in DNA glycosylase activity was observed in *NTHL1*-deficient cells specifically for 5-ohU-containing oligonucleotides, while activity toward U-containing oligonucleotides remained unaffected (Fig.1B). Additionally, trapping assay showed selective trapping of 5-ohU-containing DNA oligonucleotides in WT cell extracts, whereas this trapping was dramatically reduced in knockout cell extracts. Although we cannot exclude that other DNA glycosylases also contribute to the trapping observed, the substantial reduction in trapping activity in the *NTHL1^-/-^* cells, indicates a functional difference between WT and knockout cells (Fig.1C).

**Figure 1:**
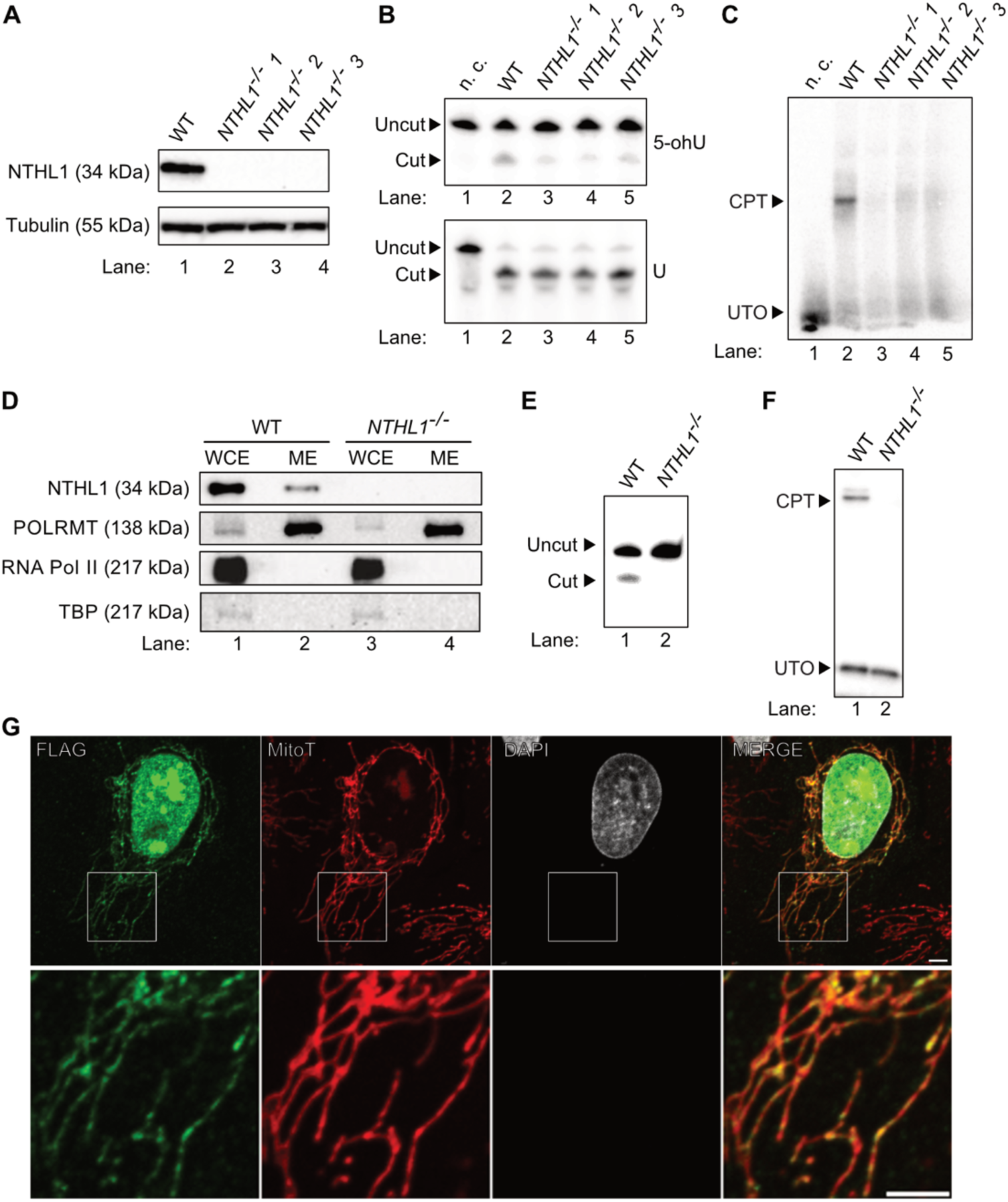
NTHL1 is active in mitochondria. **(A)** Immunoblot analysis confirming the successful knockout of *NTHL1* in HEK293 cells. Lane 1 represents WT cells, and lanes 2-4 represent independent *NTHL1^-/-^* clones. **(B)** DNA glycosylase activity in 10 μg total extracts of HEK293 WT (lane 1) and three independent HEK293 *NTHL1^-/-^* clones (lanes 2-4) on ^32^P-end-labelled double-stranded 5-ohU (upper panel) and U (lower panel). n.c.: negative control, no extract. **(C)** Covalent protein–DNA intermediates by NaCNBH3 reduction in 10 μg total extracts of HEK293 WT and three independent HEK293 *NTHL1^-/-^* clones on ^32^P-end-labelled double-stranded 5-ohU. CPT: Covalent protein-DNA intermediate; UTO: untrapped oligo. **(D)** Immunoblot analysis of HEK293 WT (lanes 1-2) and *NTHL1^-/-^* (lanes 3-4) whole cell lysates (lanes 1, 3) and mitochondrial lysates (lanes 2, 4). WCE: whole cell extract; ME: Mitochondrial extract. **(E)** DNA glycosylase activity in 10 μg mitochondrial extracts of HEK293 WT (lane 1) and HEK293 *NTHL1^-/-^* (lane 2) on ^32^P-end-labelled double-stranded 5-ohU. **(F)** Covalent protein–DNA intermediates by NaCNBH3 reduction in 10 μg mitochondrial extracts of HEK293 WT and HEK293 *NTHL1^-/-^* on ^32^P-end-labelled double-stranded 5-ohU. **(G)** Immunofluorescence analysis of nuclear and mitochondrial localization of NTHL1. U2OS cells were transfected with a plasmid coding for NTHL1-FLAG. 24 hours after transfection, mitochondria were stained with Mitotracker Red (red), fixed and stained with an antibody against FLAG (green). Nuclei were stained with DAPI (gray). Scale bar 5 µm.

To verify the mitochondrial localization and activity of NTHL1 protein, we isolated mitochondria from HEK293 cells. Immunoblotting revealed that a small fraction of NTHL1 localizes to the mitochondria (Fig.1D). To ensure signal specificity, the same experiment was performed in NTHL1-deficient cell lines, where no detectable signal was observed (Fig.1D). To assess enzymatic activity, DNA glycosylase and trapping assays were conducted using purified mitochondrial extracts from both WT and *NTHL1^−/−^* cell lines (Fig.1E, F). These assays confirmed NTHL1 enzymatic activity exclusively in WT mitochondrial extracts, validating mitochondrial isolation and functional presence of NTHL1 in mitochondria. To further confirm the mitochondrial localization of NTHL1, immunofluorescence (IF) analysis was performed and a clear colocalization of NTHL1 with mitochondrial markers was observed (Fig.1G).

### RNA Sequencing Analysis Reveals Altered Mitochondrial Gene Expression in *NTHL1^-/-^* Cell Lines

To identify whether loss of NTHL1 has an impact on the transcript levels of genes involved in mitochondrial dynamics, we performed RNA sequencing (RNA-seq) on WT and *NTHL1*^-/-^ cell lines. Differentially expressed genes (DEGs) were identified by comparative transcriptomics, with a log2 fold change threshold of +/-0.5 and false discovery rate (FDR) < 0.05. After filtering using MitoCarta 3.0, 68 mitochondrial-related genes were found to be differentially expressed in *NTHL1^-/-^* cell lines compared to WT (Fig.2A). Subsequent gene ontology (GO) enrichment analysis performed using g:Profiler ^36^ indicated that most DEGs in *NTHL1^-/-^* cell lines were associated with oxidoreductase activity as a molecular function (adjusted p-value 6.329×10^-19^) (Fig.2B). The DEGs were also enriched for biological processes such as the small molecule metabolic process and the respiratory ETC (adjusted p-values 6.329×10^-19^ and 1.660×10^-17^, respectively) (Fig.2B and fig.S2A, B). The RNA sequencing results were confirmed via RT-qPCR of genes belonging to the top identified biological processes, specifically those showing increased in mRNA levels for mitochondrial-encoded genes *MT-ND6, MT-CYTB* (Fig.S3A-B) and nuclear-encoded gene*s COX4l1, COX5B, NDUFS6, HCCS, APOOL, NFU1S, MAOA, ACSM3, ACSL6, HMOX,1 TSPO, OXCT1* (Fig.S3 C-N) as well as nuclear-encoded gene *FASN* showing decreased mRNA levels (Fig.S3O). To investigate the impact of NTHL1 deficiency on transcript levels of mitochondrial-related genes, we performed orthogonal partial least squares discriminant analysis (OPLS-DA) on mitochondrial DEGs in *NTHL1^-/-^* and WT samples. The OPLS-DA score plot (Fig.S2C) revealed a clear separation between KO and WT samples along the first predictive component, indicating distinct mitochondrial gene expression profiles between the groups. The clustering of biological replicates within each condition highlights the consistency of transcriptional changes in response to NTHL1 deficiency. The corresponding loadings plot (Fig.S2D) identified genes contributing to the observed separation. Genes associated with KO samples, such as *ND6, ND3, ND4L* are involved in mitochondrial ETC function, reflecting mitochondrial respiratory activity. Genes associated with WT samples, including *FASN* and *ALDH1L2*, represent metabolic functions such as lipid biosynthesis and one-carbon metabolism. These findings suggest that NTHL1 deficiency alters mitochondrial gene expression, potentially leading to functional reorganization of mitochondrial processes. We used DModX residual analysis to evaluate potential outliers, and all samples falling below the critical threshold (DCrit = 0.05), confirming the robustness of the analysis (Fig.S2E).

**Figure 2:**
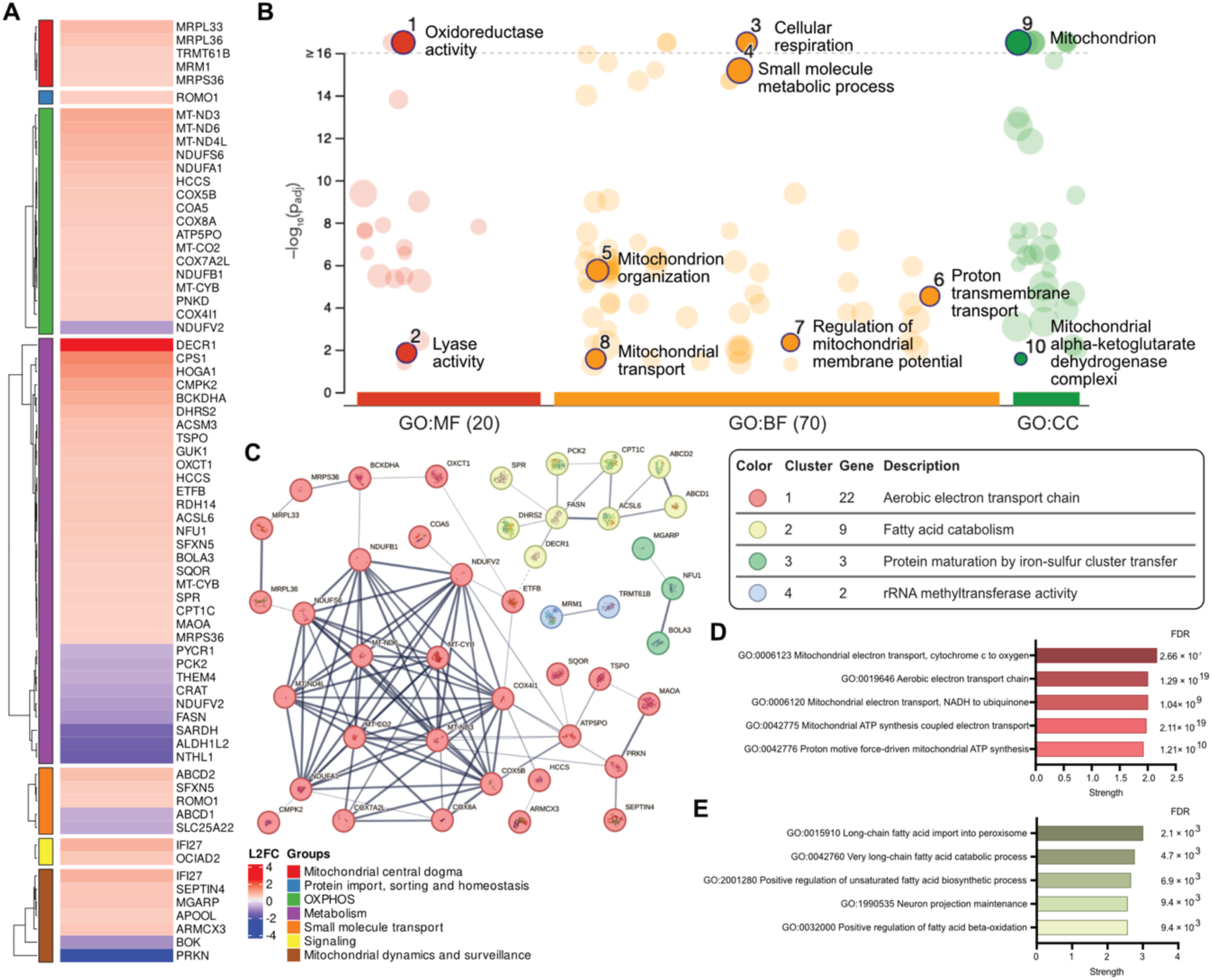
Impact of *NTHL1* knockout on gene expression of mitochondrial-related genes in HEK293 cells. **(A)** Heatmap showing DEGs identified by RNA-sequencing (RNA-seq) in *NTHL1^-/-^*. Red colored boxes indicate significantly increased mRNA levels, and blue boxes indicate significantly diminished mRNA levels in *NTHL1^-/-^* cell lines compared to WT (log2 fold change threshold +/-0.5, FDR < 0.05). Analysis was performed in two independent *NTHL1^-/-^* clones. Legend is in the bottom right. **(B)** GO Enrichment Analysis: The x-axis represents the GO term (biological process, molecular function, cellular component), and the y-axis represents adjusted p-value (-log10 Padj). Top significant driver terms are highlighted (adjusted p-value < 0.05). (**C-E**) Network analysis and Gene Ontology (GO) Analysis of mitochondrial-related genes altered in *NTHL1^-/-^* cell Lines. **(C)** STRING network analysis of mitochondrial DEGs in *NTHL1^-/-^* cell lines. Network is subdivided in four clusters (depicted in red, light green, blue and dark green) through k-means clustering, with the members that characterize each cluster presented. Edge thickness is representative of interaction confidence. **(D)** Gene Ontology (GO) enrichment analysis for biological processes (represented with strength as a scale) shows significant enrichment for terms related to aerobic ETC and oxidoreductase activity in Cluster 1. **(E)** GO analysis of biological processes related to Cluster 2 highlights significant enrichment for processes involved in fatty acid metabolism, lipid catabolism, and arachidonate-CoA ligase activity, with strength represented on a scale. FDR: false discovery rate.

To further explore the functional relationships among the mitochondrial-related genes affected by NTHL1 loss, we performed a STRING network analysis to identify key interaction networks among the differentially expressed mitochondrial-related genes (Fig.2C). Four distinct clusters emerged, with cluster 1 being the largest, comprising 29 genes and was primarily associated with mitochondrial functions crucial for energy production (Fig.2D). This cluster was significantly enriched for genes involved in the aerobic ETC and oxidoreduction-driven active transmembrane transporter activity, both critical for mitochondrial respiration and ATP synthesis. Cluster 2 (gene count: 9) included genes involved in fatty acid catabolism and arachidonate-CoA ligase activity, which are essential for mitochondrial lipid metabolism (Fig.2E). In addition to these main clusters, two smaller clusters were identified: Cluster 3 (gene count: 3) was associated with protein maturation by iron-sulfur cluster transfer, and Cluster 4 (gene count: 2) was linked to rRNA methyltransferase activity, a process crucial for mitochondrial protein synthesis. Alterations in this cluster point to potential defects in mitochondrial translation and protein maturation in *NTHL1^-/-^* cells. Hence, NTHL1 deficiency affects multiple mitochondrial processes, including energy production and fatty acid metabolism.

### NTHL1 deficiency causes increased mtDNA-CN and promotes mitochondrial biogenesis and fusion

Among the DEGs several mitochondrially encoded genes displayed increased levels. Given that mitochondrial gene expression is tightly linked to mitochondrial function, we reasoned that this overall increase in mRNA levels might be a compensatory response to mitochondrial dysfunction, potentially driven by an increase in mitochondrial biogenesis or a response to elevated oxidative stress. mtDNA copy number (mtDNA-CN) has long been recognized as a biomarker for mitochondrial dysfunction^37^. Analysis of mtDNA-CN revealed a 2-fold increase in *NTHL1^-/-^* compared to WT cell lines (Fig.3A). To assess mtDNA integrity, we used two qPCR-based methods to quantify DNA damage. First, the Real-time qPCR (RT-qPCR) Analysis of Damage Frequency (RADF) assay^38^, which measures DNA lesions that block TaqI enzyme digestion at 5’-TCGA-3’ sites, revealed increased mtDNA damage in NTHL1-deficient cells (Fig.3B). Similarly, long-amp qPCR, which detects DNA damage by measuring reduced amplification of long mtDNA fragments, showed significantly lower amplification in NTHL1-deficient cells compared to WT (Fig.3C). This reduction reflects impaired polymerase processivity due to DNA damage, indicating increased proportion of lesions per mtDNA molecule. Together, these assays consistently indicate elevated mtDNA damage upon loss of *NTHL1*.

**Figure 3:**
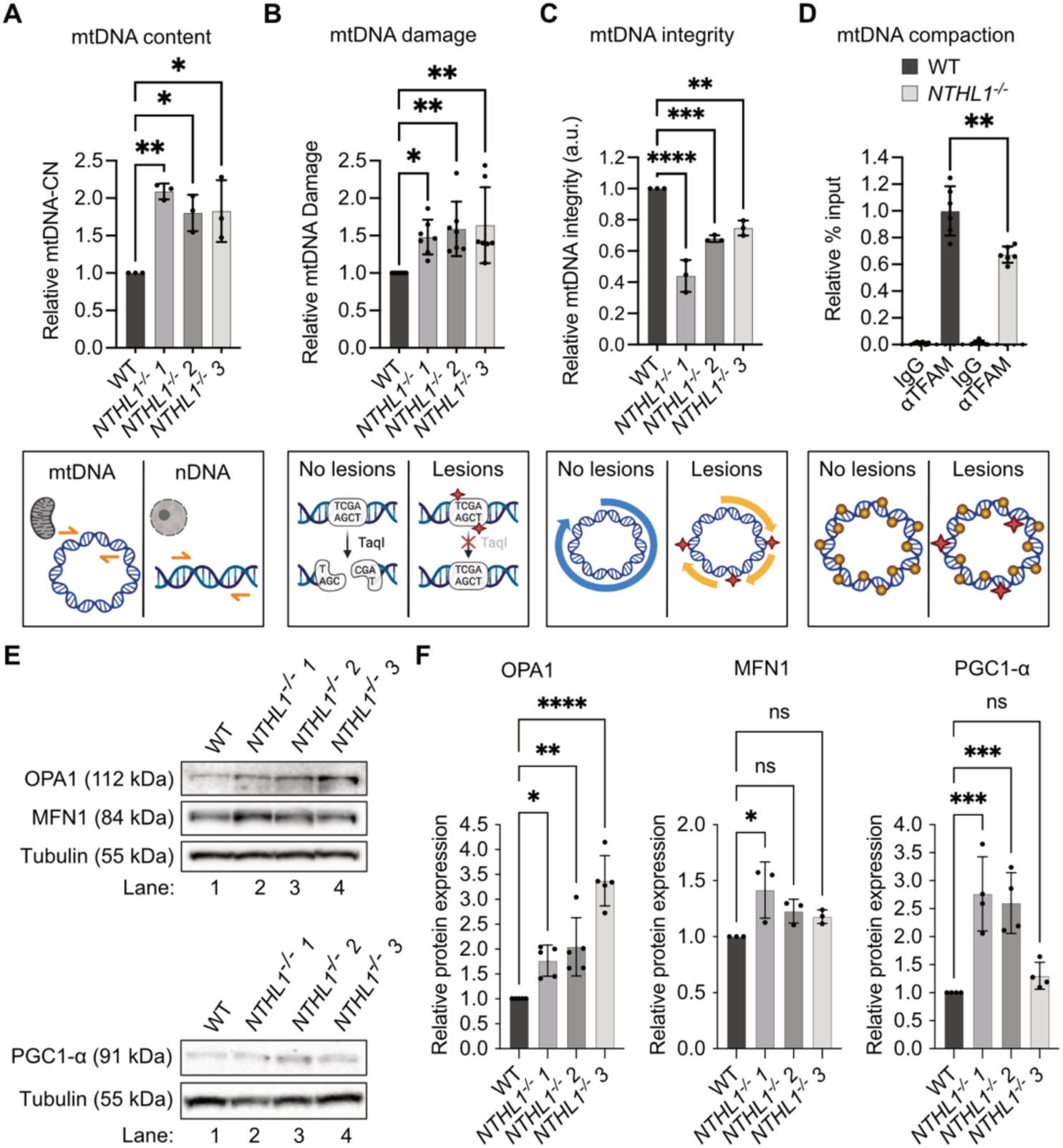
*NTHL1^-/-^* cells show increased mitochondrial biogenesis. **(A)** Relative mtDNA copy number of HEK293 WT and three independent HEK293 *NTHL1^-/-^* clones**. (B)** Relative mtDNA damage measured with the Real-time qPCR Analysis of Damage Frequency (RADF) assay of HEK293 WT and three independent HEK293 *NTHL1^-/-^* clones**. (C)** Relative mtDNA damage measure via long-amplicon quantitative in HEK293 WT and three independent HEK293 *NTHL1^-/-^* clones**. (D)** ChIP on mtDNA for TFAM in HEK293 WT and HEK293 *NTHL1^-/-^* cells. **(A-D)** Below each bar graph is a graphical representation showing the method used to perform the assay. (**E**) Immunoblot analysis of OPA1, MFN1, and PGC1-α protein levels in WT and *NTHL1^-/-^* cell extracts. **(F)** Relative quantification of data depicted in E. Error bars represent mean ± SD (n ≥ 3). *p ≤ 0.05; **p ≤ 0.01; ***p ≤ 0.001; ****p ≤ 0.0001, one-way ANOVA.

To evaluate whether DNA lesions affect mtDNA compaction we performed chromatin immunoprecipitation (ChIP) targeting TFAM, a mtDNA coating and compacting protein^39,40^. Our analysis revealed a significant reduction in TFAM binding in *NTHL1*^-/-^ cell lines compared to WT cells (Fig.3D). This decrease was not attributed to altered TFAM expression levels, as RT-qPCR and immunoblotting analyses showed no changes in expression levels between WT and *NTHL1^-/-^* cell lines (Fig.S3A-C).

Next, to determine whether mitochondrial dynamics are further affected, we performed immunoblotting for mitochondrial fusion proteins. We observed an average 2-fold increase in the protein levels of Dynamin-like GTPase OPA1 (Fig.3A, B), a mitochondrial inner membrane protein that plays a central role in inner membrane fusion and cristae morphology^41–45^. Notably, loss of OPA1 disrupts respiratory chain supercomplex assembly, leading to reduced electron transport chain (ETC) activity and impaired OXPHOS^46^. Furthermore, knockout of OPA1 in fibroblasts has previously been shown to result in decreased mtDNA levels^47^.

Furthermore, our results showed no significant differences in the protein levels of either Mitofusin 1 (MFN1) and Mitofusin 2 (MFN2)— other key mitochondrial outer membrane proteins that mediate mitochondrial fusion— between *NTHL1^-/-^* and WT cells (Fig.3A, B). We also observed an average 2-fold increase in the protein levels of peroxisome proliferator-activated receptor γ coactivator 1α (PGC1α), a key regulator of mitochondrial biogenesis. Under oxidative stress, PGC1α drives the expression of numerous nuclear-encoded mitochondrial genes, including those involved in mitochondrial biogenesis and oxidative phosphorylation, aligning with our transcriptomics results. Taken together with the increase in mtDNA-CN, our results indicate that NTHL1 plays a role in regulating mitochondrial biogenesis, potentially mediated by PGC1α. Furthermore, the increase in OPA1 levels suggests that loss of NTHL1 enhances mitochondrial fusion.

### Loss of NTHL1 is associated with increased mitochondrial mass and respiration

To further understand the role of NTHL1 in regulating mitochondrial function, we first assessed oxidative phosphorylation (OXPHOS) protein levels and observed a general increase in NTHL1 deficient cells (Fig.4A, B). We also performed IF using MitoView green to quantify mitochondrial content and observed a marked increase in mitochondrial staining in *NTHL1*^-/-^ cell lines (Fig.4C, D). Transmission electron microscopy revealed changes in mitochondrial morphology with *NTHL1^-/-^* cells showing an increased mitochondrial area (Fig.4E, F).

**Figure 4:**
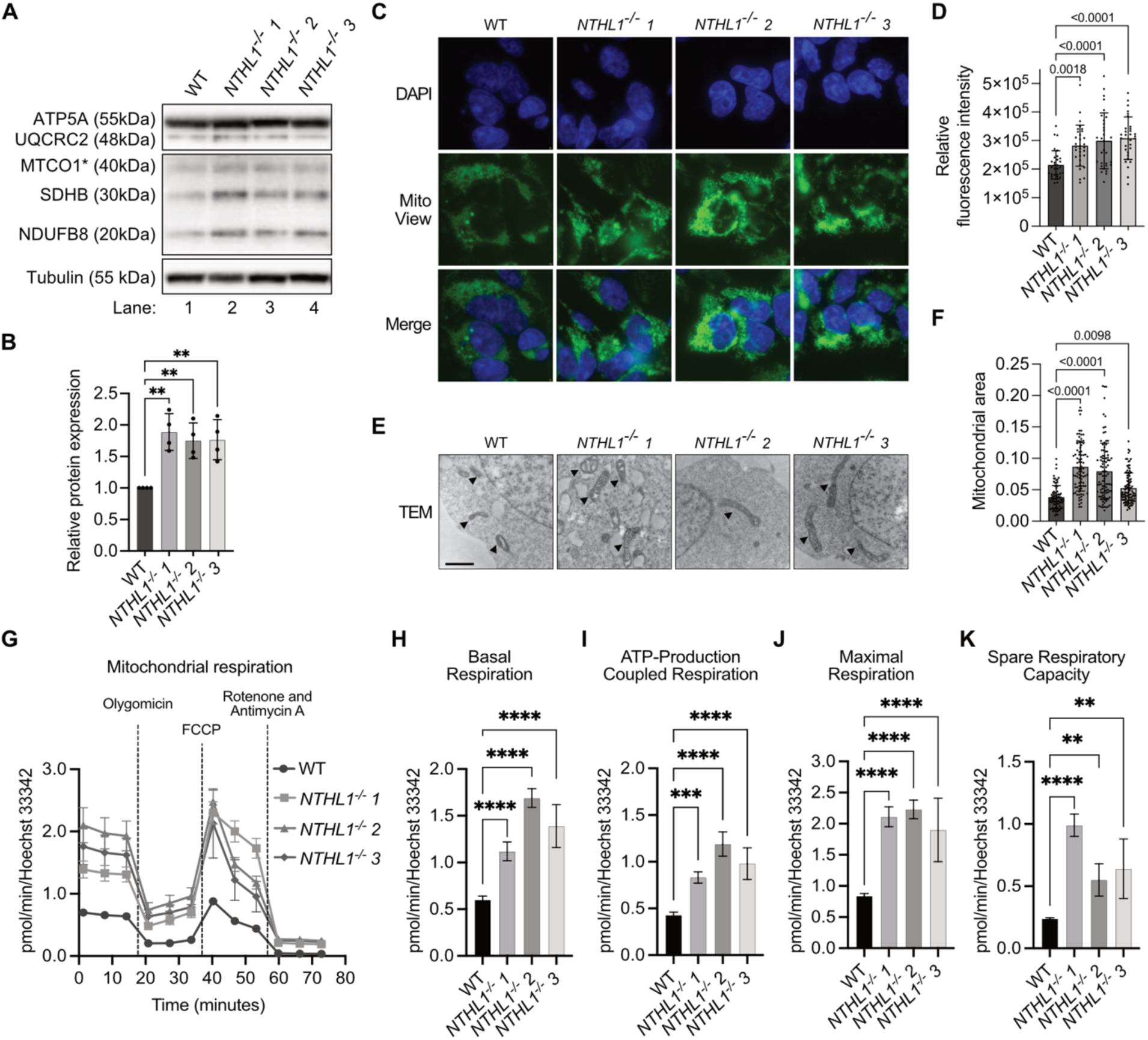
Mitochondrial mass and energy metabolism is enhanced *NTHL1^-/-^*cells. **(A)** Immunoblot analysis of OXPHOS complex proteins of HEK293 WT (lane 1) and three independent HEK293 *NTHL1^-/-^* clones (lanes 2-4). CV-ATP5A, CIII-UQCRC2, CIV-MTCO1*, CII-SDHB, CI-NDUFB8. *Mitochondrially encoded **(B)** Quantification of total OXPHOS protein levels from immunoblots. **(C)** Representative immunofluorescence images of WT and three independent HEK293 *NTHL1^-/-^* clones. Cells were stained using MitoView (green), and DAPI for nuclear visualization (blue). Merged images show increase mitochondria signal intensity in *NTHL1^-/-^* lines. **(D)** Quantification of mitochondrial mass of the experiment depicted in C. **(E)** Representative transmission electron microscopy (TEM) images of WT and *NTHL1^-/-^* cells. Scale bar = 2 μm. Arrows indicate single mitochondria. **(F)** Quantification of mitochondrial size from TEM images. **(G-K)** mitochondrial oxygen consumption curves are presented as averages ± standard deviations for each measurement time point (n = 5). The initial three measurement points represent basal mitochondrial respiration. The following three points, after the addition of oligomycin to block adenine nucleotide translocation, represent proton leak-stimulated oxygen consumption. Next, three points represent maximal mitochondrial oxygen consumption capacity, achieved by uncoupling the mitochondrial inner membrane with FCCP. The final three points measure non-mitochondrial oxygen consumption when the mitochondrial respiratory chain is inhibited by rotenone and antimycin A. **(G)** Summary data for basal respiration. **(H)** Summary data of ATP-Production (**I**) coupled respiration, (**J**) maximal respiration, **(K)** spare respiratory capacity. Error bars represent mean ± SD (n ≥ 3). *p ≤ 0.05; **p ≤ 0.01; ***p ≤ 0.001; ****p ≤ 0.0001, one-way ANOVA.

To examine the impact of increased mitochondrial mass in *NTHL1^-/-^* cells, we assessed mitochondrial metabolic activity using the Seahorse XF Cell Mito Stress Assay (Fig.4G-K). The average basal mitochondrial oxygen consumption rate (OCR) in *NTHL1^-/-^* cell lines was on average 80% higher than in WT cells (Fig.4G, H). This positive trend was also evident for mitochondrial OCR sensitive to the ATPase inhibitor oligomycin, where the OCR coupled to ATP production was significantly elevated in *NTHL1^-/-^* cells compared to WT cells (see OCR after addition of oligomycin in Fig.4G, and bars in Fig.4I). Moreover, when mitochondria were stressed by permeabilizing the inner membrane for H^+^ with carbonyl cyanide 4-(trifluoromethoxy) phenylhydrazone (FCCP) to reveal maximal mitochondrial respiratory capacity, the responsive increase in OCR of *NTHL1^-/-^* cell mitochondria were on average 2 times higher compared to WT cells (see OCR after addition of FCCP in Fig.4G and bars in Fig.4J). Additionally, the spare respiratory capacity was significantly increased in *NTHL1^-/-^* cells (see OCR after addition of Rotenone and Antimycin in Fig.4G and bars in Fig.4K). These findings suggest that the absence of NTHL1 enhances mitochondrial respiratory activity and overall mitochondrial efficiency. This enhancement suggests, that while NTHL1 is essential for maintaining mtDNA integrity, its absence triggers adaptive mechanisms to sustain mitochondrial function and energy production. Thus, the phenotype could be a compensatory response to increased mtDNA damage observed in *NTHL1^-/-^* cells.

Furthermore, the Seahorse analysis revealed that NTHL1 deficient lines showed increased proton leak and non-mitochondrial oxygen consumption (Fig.S5A, B). Increased proton leak suggests that more protons (H⁺) are crossing the mitochondrial membrane independently of ATP production resulting in a “leak” of energy as heat, bypassing the ATP synthase. Non-mitochondrial oxygen consumption reflects the oxygen consumed by reactions outside the mitochondria. These reactions could include oxidase activities, such as NADPH oxidases or other enzymes that generate ROS indicating heightened oxidative stress.

Taken together, these observations indicate that the cells are experiencing mild oxidative stress, which may be driving adaptive changes in cellular metabolism and energy production. This shift is reminiscent of mitohormesis—a compensatory response where moderate mitochondrial stress enhances cellular resilience and function. This interpretation aligns with previous observations showing that NTH-1 loss enhances resilience in a *C. elegans* model^10^.

### Loss of NTHL1 reduces sensitivity to Complex I inhibition

If the changes observed were indeed part of a mitohormetic response, we would expect that the *NTHL1^-/-^* cells would exhibit altered response to mitochondrial stress due to the observed increase in OXPHOS-related proteins in these cells. MPP+ is a mitochondrial toxin known to inhibit mitochondrial complex I (NADH oxidoreductase) of the ETC ^48–50^. MPP+ disrupts electron flow by inhibiting complex I, reducing ATP production and increasing ROS, leading to oxidative stress and cellular damage. We hypothesized that NTHL1 deficiency might alter the cellular response to MPP^+^-induced mitochondrial dysfunction. Consistent with our hypothesis, NTHL1 deficient cells exhibited enhanced tolerance to MPP+ compared to WT cells. After 72 hours of MPP+ treatment, WT cells showed a significant reduction in cell numbers, while *NTHL1^-/-^* cells maintained stable cell counts across various MPP+ concentrations (Fig.5A). Furthermore, cell viability was decreased in both WT and *NTHL1^-/-^* cells following MPP+ treatment, but *NTHL1^-/-^* cells maintained significantly higher viability than WT cells at all tested concentrations (Fig.5B). Similar resistance was observed in HAP1 cell lines (Fig.S5A-C).

**Figure 5:**
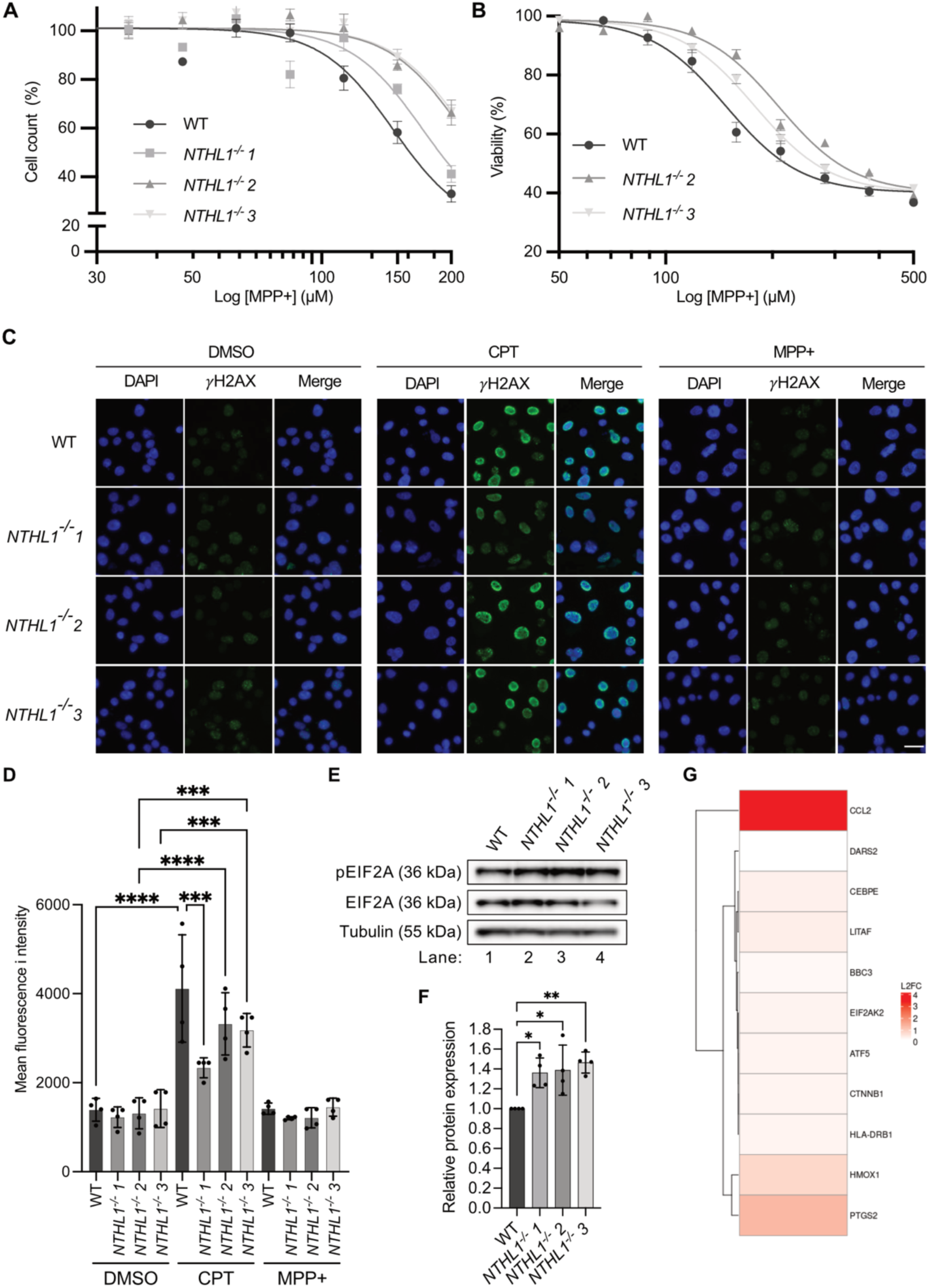
NTHL1-deficient cells display enhanced tolerance to 1-methyl-4 phenylpyridinium (MPP+). **(A)** HEK293 WT and three independent HEK293 *NTHL1^-/-^* clones were treated with the indicated concentrations of MPP⁺ for 72 h, stained with Hoechst 33342, and the number of cells per well was counted. MPP⁺ treatment reduced cell numbers in WT but not in *NTHL1^-/-^* cells. **(B)** MPP+ treatment decreases viability. WT and *NTHL1^-/-^* cells were exposed to the indicated concentrations of MPP+ for 72 h, and viability was assessed with Cell-titer glo. (**C**) Representative immunofluorescence image of γH2AX in WT and *NTHL1^-/-^* cells following CPT or MPP+ treatment. **(D)** Quantification of the analysis depicted in C. **(E)** Immunoblot analysis of pEIF2A and EIF2A in extracts from HEK293 WT (lane 1) and three independent HEK293 *NTHL1^-/-^* clones (lanes 2-4). **(F)** Quantification of the data shown in panel F. **(G)** ATF4-regulated genes are activated in *NTHL1^-/-^* cell lines. Error bars represent mean ± SEM (A, B, D) or ± SD (n ≥ 3) (G), representative of n=2 (A, B, D) or n=5 independent experiments (G). *p ≤ 0.05; ***p ≤ 0.001; ****p ≤ 0.0001, one-way ANOVA.

We reasoned that this effect was primarily mitochondrial and that MPP+ did not trigger a nuclear DNA damage response. To confirm this, we treated cells with MPP+ and performed IF staining for γH2AX, a marker of DNA damage. No induction of γH2AX foci were observed in WT or NTHL1^−/−^ cells upon MPP+ treatment (Fig.5C, D). As a control, we treated cells with Camptothecin (CPT), a topoisomerase I inhibitor that induces DNA damage by stabilizing the cleavage complex between topoisomerase I and DNA, leading to replication-associated double-strand breaks. As expected, CPT treatment led to a significant increase in γH2AX foci (Fig.5C, D). These findings support the idea that loss of NTHL1 induces pathways involved in mitochondrial stress response and cell proliferation under conditions of complex I inhibition, further reinforcing that NTHL1 deficiency specifically influences mitochondrial stress responses.

### Loss of NTHL1 activates the integrated stress response

To explore how NTHL1 deficiency affects cellular stress responses, we investigated eIF2α phosphorylation levels in *NTHL1^-/-^* cells, as it plays a critical role in the integrated stress response (ISR). Immunoblotting confirmed increased eIF2α phosphorylation in *NTHL1^-/-^* cells compared to WT (Fig. 5E, F), indicating activation of the ISR pathway. Given the central role of ATF4 as a downstream effector of phosphorylated eIF2α and its rule of mitochondrial stress response^51^, we next examined the expression of ATF4 target genes. Several ATF4-regulated genes were upregulated in *NTHL1^-/^*^-^ cells, including ATF5, BBC3, and CCL2. ATF5 is involved in cellular stress responses and mitochondrial maintenance^52,53^, BBC3 (also known as PUMA) promotes apoptosis^54^, and CCL2 is linked to metabolic resonse^55^. Other ATF4 targets that showed increased expression included CEBPE, CTNNB1 and DARS2, which encodes a mitochondrial aspartyl-tRNA synthetase essential for mitochondrial protein synthesis. We also observed increased expression of EIF2AK2, a kinase that phosphorylates eIF2α and amplifies the ISR, suggesting a positive feedback loop reinforcing stress adaptation. Furthermore, genes involved in oxidative stress responses and inflammation, such as HMOX1 LITAF, and PTGS2 were also upregulated.

The observed upregulation of ATF4 target genes, suggests that the ISR is at least partially actively engaged in *NTHL1^-/-^* cells as adaptive mechanisms to counteract mitochondrial dysfunction and oxidative stress. This is further supported by their increased resistance to mitochondrial stress (e.g., MPP+ treatment), aligning with the principles of mitohormesis.

## DISCUSSION

Here, we shed new light on the interplay between DNA repair and mitochondrial regulation, an area that remains poorly understood. Although NTHL1 has been previously shown to localize in mitochondria^56^, its functional role in this organelle remained unclear. Our study shows that NTHL1 deficiency alters the transcript levels of mitochondrial-related genes and triggers a mitochondrial stress response. This finding parallels with previous studies showing that DNA glycosylases can influence transcription in the nucleus^35,57,58^. Despite the expected accumulation of mitochondrial DNA damage in NTHL1-deficient cells (Fig.3A-D), we unexpectedly observed an increase in mtDNA copy number, uncovering a complex interplay between DNA lesion accumulation and mitochondrial DNA content. This increase suggests a compensatory mechanism that challenges the canonical view that mitochondrial genome instability inevitably leads to genome depletion^6,59–61^. It also raises new questions regarding the regulatory balance between DNA damage accumulation and mitochondrial biogenesis.

Although causality remains to be determined, the observed mtDNA damage accompanied by reduced TFAM binding suggests that specific DNA lesions hinder TFAM’s interaction with mtDNA. The TFAM/mtDNA ratio regulates the compaction and accessibility of mtDNA both *in vitro*^62^ and in cells ^63^. Reduced TFAM /mtDNA ratio could allow for relaxation of the nucleoids, permitting increased replication. Additionally, TFAM is unevenly distributed along mtDNA, with distinct high-affinity binding sites^39^, a pattern likely influenced by multiple factors, including TFAM–TFAM interactions^64^, cross-strand binding^62^, and sequence-specific modulation of TFAM-DNA binding^65^. While TFAM has been shown to process AP sites^66^ and to modulate the activity of monofunctional DNA glycosylases such as UNG and OGG1^67^, its potential interaction with bifunctional DNA glycosylases like NTHL1 remains unclear. Given that bifunctional DNA glycosylases can process AP sites independently, TFAM and NTHL1 may have opposing roles in mtDNA maintenance.

NTHL1-deficient cells exhibited increased mitochondrial mass, fusion dynamics (Fig.3, 4), and enhanced respiratory efficiency (Fig.4), suggesting a previously unrecognized link between DNA lesion accumulation and mitochondrial network remodeling. These findings align with previous studies using cell models with stably reduced mtDNA-CN, which correlated with decreased mitochondrial protein expression and a decline in respiratory enzyme activity^68^. Furthermore, *NTHL1*^-/-^ cells exhibited increased tolerance to mitochondrial stress, particularly in response to Complex I NADH oxidoreductase inhibition by MPP+^45,48–50^ (Fig.5A, B) revealing a beneficial functional adaptation. Although no DNA glycosylase deficiency has been previously linked to MPP+ resistance in humans, Sanders *et al.* demonstrated that genetic variants of the BER genes increase risk of PD in combination with pesticide exposures, which are known to affect mitochondrial function^69^. Furthermore our results align with observation in a *C.elegans* PD model in which *nth-1* mutants exhibit similar resistance to mitochondrial stressors^10^.

Our findings suggest that loss of NTHL1 may trigger compensatory mechanisms aimed at sustaining mitochondrial function in the face of increased oxidative stress. Despite these adaptive changes, the observed increase in proton leak and non-mitochondrial oxygen consumption points toward heightened oxidative stress and the potential involvement of ROS-generating pathways.

Collectively, our findings align with the concept of mitohormesis, where mild mitochondrial stress can induce adaptive cellular responses that improve resilience. Additionally, mitochondrial stress does not operate in isolation but influences cytosolic proteostasis networks, modulating protein synthesis, stability, and degradation to maintain cellular metabolism and proliferation. Our findings suggest that NTHL1 deficiency may reshape these networks, shifting the balance toward stress resilience rather than dysfunction. This broader perspective on mitochondrial stress responses underscores the intricate crosstalk between DNA repair, mitochondrial homeostasis and cellular adaptation, with potential implications for neurodegeneration, aging, and cancer. The increase in mitochondrial biogenesis-related genes and the activation of stress response pathways align with findings from other studies in the field which show that cells often respond to mitochondrial dysfunction by enhancing biogenesis and mitochondrial protein synthesis^70^. Additionally, the interaction of NTHL1 with the ISR and mitohormesis pathways introduces a new perspective on how mitochondrial stress is managed at the cellular level.

While the mechanistic basis of nuclear DNA repair defects triggering stress response through interference with the transcription machinery is well understood, the underlying processes in mitochondria remain unclear. A reduction in mtDNA compaction suggests that mtDNA may adopt a more open conformation, potentially facilitating increased rates of replication and/or transcription. Future research should focus on determining the causality between mtDNA damage and compensatory mitochondrial adaptations. Specifically, it should investigate whether DNA lesions interfere with the binding of regulatory proteins such as TFAM causing changes in replication or metabolic processes. Alternatively, it should explore if increased mtDNA copy number, without a matched increase in TFAM levels, contributes to the observed phenotype.

Additionally, pinpointing the specific DNA lesions that accumulate in *NTHL1^-/-^* cells is crucial, as they may drive the observed mitochondrial phenotype and establish a direct link between DNA repair deficiency and its functional consequences on cellular health.

Another critical aspect is understanding the long-term consequences of NTHL1 deficiency. Over time, observed compensatory mechanisms might lead to a state of metabolic imbalance or oxidative stress, potentially contributing to cellular aging or other dysfunctions. To establish a connection to age-related diseases, future research should assess the impact of NTHL1 on cellular lifespan and metabolic plasticity using human aging models.

Overall, our findings open new directions for research into the interplay between mitochondrial DNA quality control and cellular stress responses. Future studies will be essential to fully elucidate NTHL1’s role in mitochondrial homeostasis and its broader impact on cellular physiology and human disease.

## ACKNOWLEDGMENTS

We extend our gratitude to Veronica Suaste Morales for her assistance with the quality control of the high-throughput data. We sincerely thank Professor Lars Eide for his valuable support and insightful discussions, and Magnar Bjørås for generously providing HAP1 cell lines.

## AUTHOR CONTRIBUTIONS

Lisa Hubers: Investigation, Formal analysis, Validation, Visualization, Writing - original draft Yohan Lefol: Data curation, Formal analysis, Investigation

Alexander Myhr Skjetne, Luisa Luna, Solveig Osnes Lund, Ane Marit Wågbø, Xavier Renaudin, Annikka Polster, Anders Knoph Berg-Eriksen, Francisco Jose Naranjo-Galindo: Investigation, Validation.

Anna Campalans and Torkild Visnes: Supervision

Hilde Loge Nilsen: Conceptualization, Funding acquisition, Supervision, Writing - original draft

Nicola Pietro Montaldo: Conceptualization, Formal analysis, Funding acquisition, Investigation, Validation, Project administration, Supervision, Visualization, Writing - original draft

Supplementary Data are available as separate pdf file at NAR online.

## DATA AVAILABILITY

Raw files (Fastq) and processed counts associated with RNA sequencing data presented in Fig.2 is available in GEO under accession number GSE291371.

---

**Table 1.** List of antibodies

| Reagent or Resource | Source | Identifier |
| --- | --- | --- |
| Goat Anti-Rabbit IgG Antibody (H+L), HRP | Vector Laboratories | PI-1000-1 |
| Horse Anti-Mouse IgG Antibody (H+L), HRP | Vector Laboratories | PI-2000-1 |
| Anti-NTH1 antibody [2660C1a] | Abcam | ab70726 |
| Anti-Alpha Tubulin antibody [DM1A], HRP | Abcam | ab40742 |
| Anti-POLRMT antibody | Abcam | ab32988 |
| Anti-RNA polymerase II CTD repeat YSPTSPS (phospho S5) antibody | Abcam | ab5131 |
| Anti-OPA1 Antibody (D-9) | Santa Cruz | sc-393296 |
| Anti-Total OXPHOS Rodent WB Antibody Cocktail | Abcam | ab110413 |
| Anti-Mitofusion 1+2 antibody (3C9) | Abcam | ab57602 |
| Anti-PGC1α antibody | Abcam | ab191838 |
| Anti-TFAM antibody | ProteinTech | 22586-1-AP |
| Anti-eIF2α antibody | Cell Signalling Technology | 9722 |
| Anti- Phospho-eIF2α antibody (Ser51) | Cell Signalling Technology | 9721 |
| Anti-p-Histone H2A.X Antibody (Ser 139) | Santa Cruz | sc-517348 |
| Goat anti-Mouse IgG (H+L) Highly Cross-Adsorbed Secondary Antibody, Alexa Fluor™ 488 | Invitrogen | A11029 |
| Anti-Flag M2 | Sigma-Aldrich | F1804 |

**Table 2.** List of kits and reagents

| Reagent or Resource | Source | Identifier |
| --- | --- | --- |
| Mitochondria isolation kit | Miltenyi | 130-094-532 |
| AllPrep DNA/RNA/Protein Mini Kit | Qiagen | 80004 |
| MitoView Green | Biotium | 70054 |
| Mitotracker Red | Invitrogen | M22425 |

**Table. 3.**
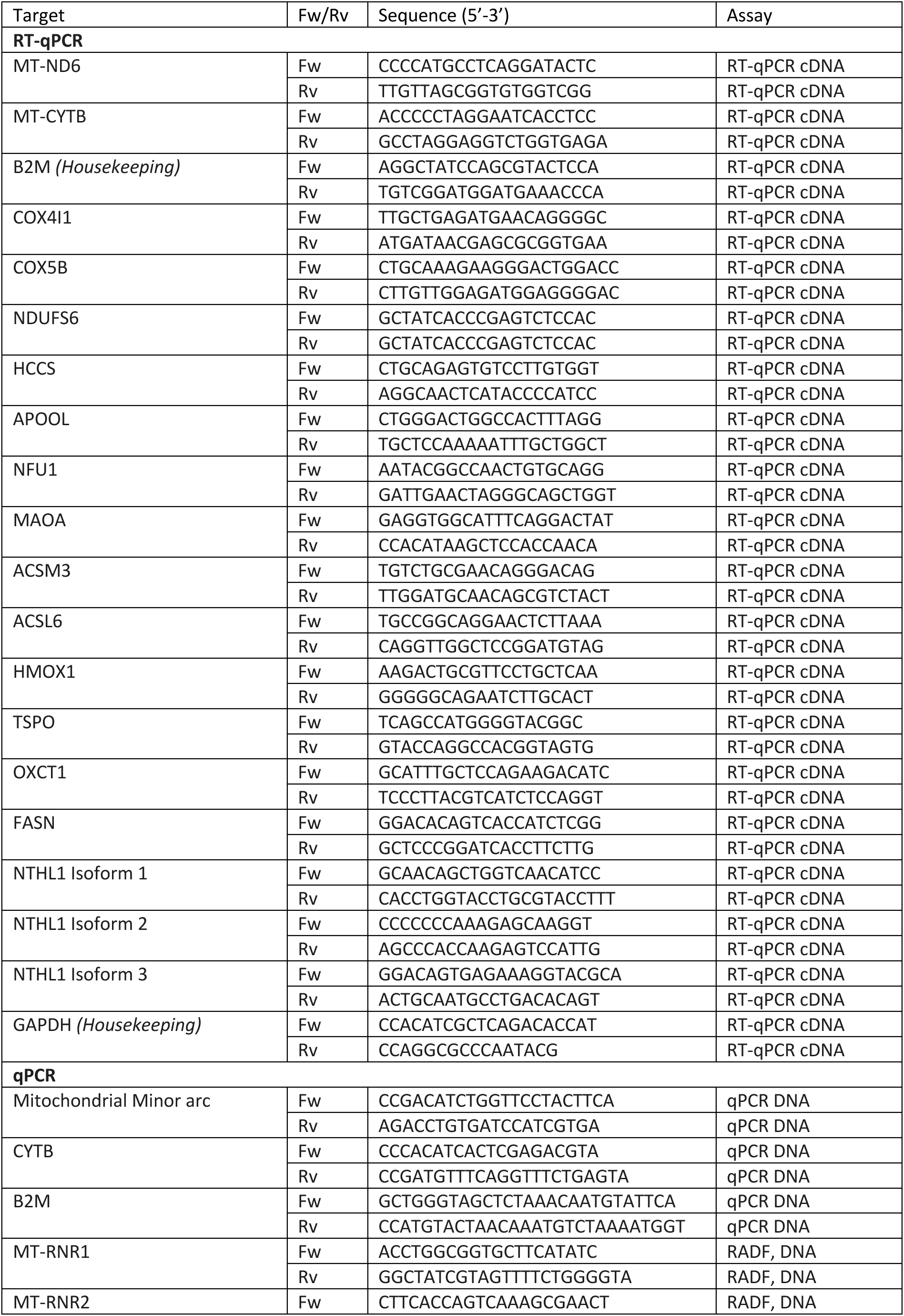
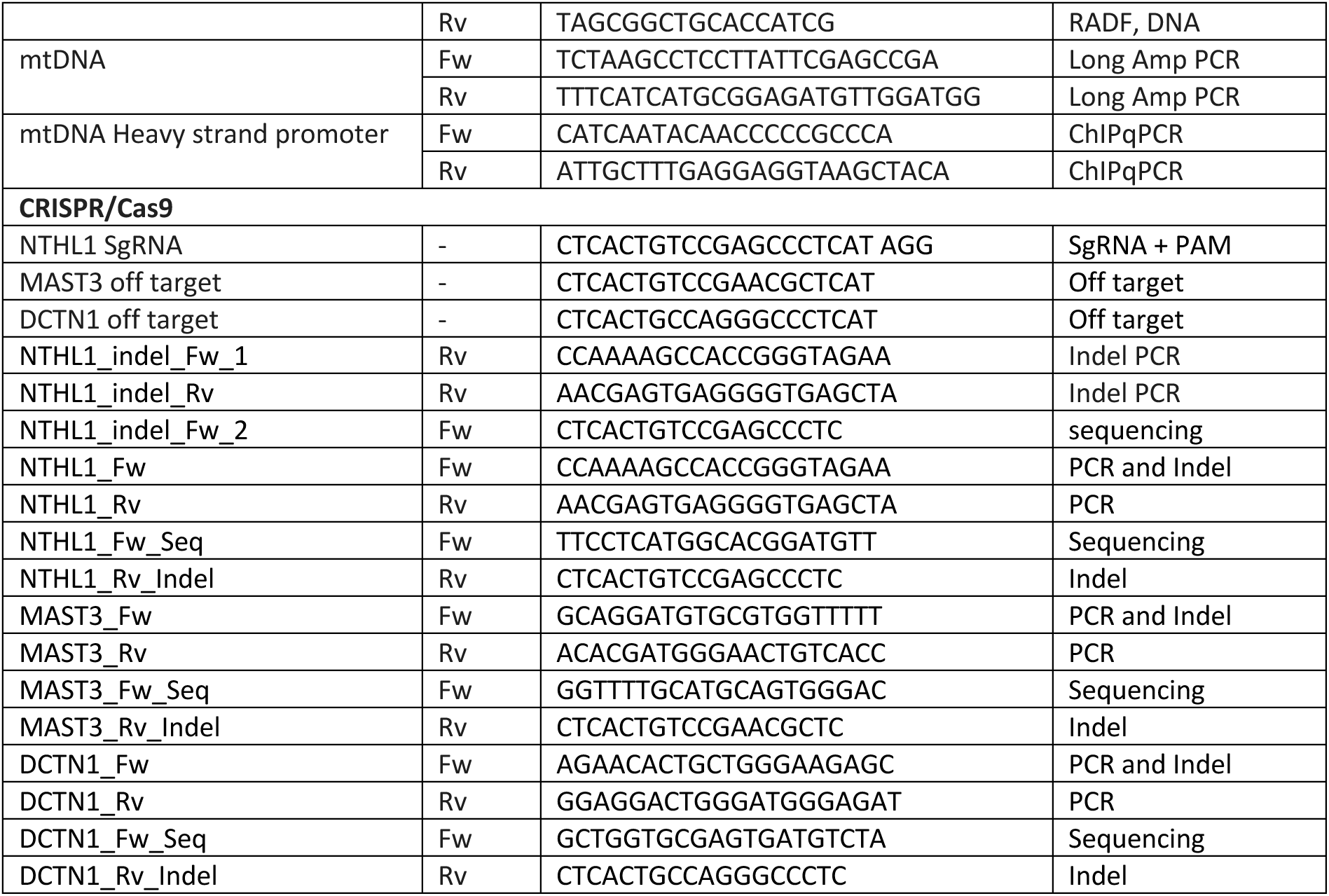
List of primers and sequences

