## Supplementary data for "Functional Impact of Nth-like DNA glycosylase on Mitochondrial Dynamics"

### Figure S1

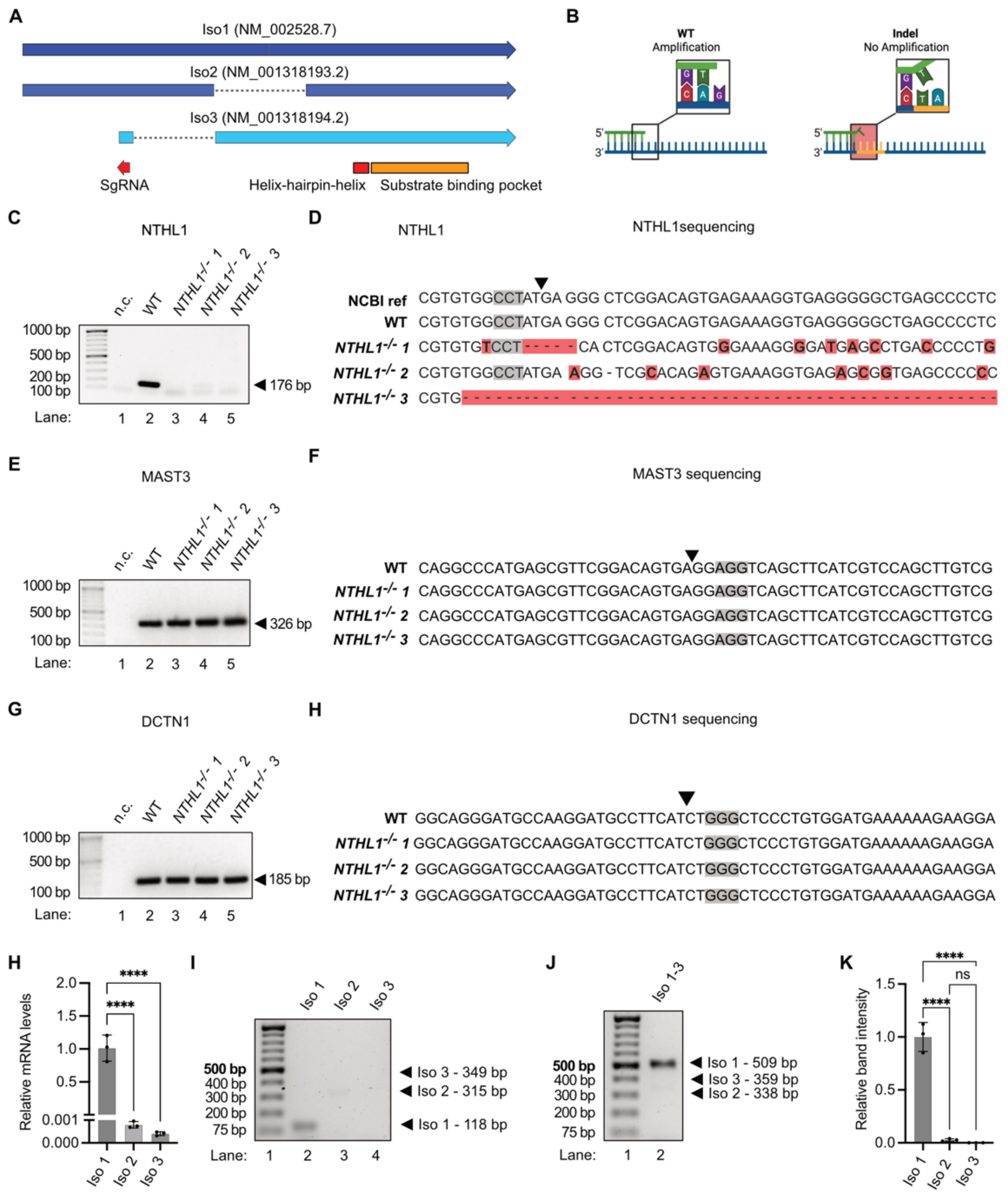


**Figure S1: Screening and quality control of *NTHL1*$^{-/-}$ clones. (A)** Schematic representation of the CRISPR-Cas9 design. The cut site (red) is shown in relation to the exons (blue) on all three isoforms of NTHL1. Since the cut is upstream of the substrate-binding pocket, the resulting protein is non-functional. **(B)** Indel-specific PCR: An indel-specific PCR was performed to screen for mutations in *NTHL1* knockout clonal cell lines. The PCR does not yield a product in the presence of indel mutations, confirming successful targeting. **(C)** Clonal screening: Individual clonal cell lines were isolated and screened for indel mutations in the sgRNA-targeted region of *NTHL1* using indel-specific PCR, confirming gene disruption at the DNA level. **(D)** Sanger sequencing validation: Indel mutations identified in selected knockout clones were further validated by Sanger sequencing, confirming the precise successful editing made by CRISPR/Cas9. **(E-H)** Off-target analysis: To ensure the specificity of the CRISPR/Cas9 system, potential off-target sites identified by CRISPOR were examined. Exonic off-target regions in MAST3 **(E)** and DCTN1 **(G)** were screened using indel-specific PCR, and the amplified regions were further confirmed to be free of mutations by Sanger sequencing **(F, G)**. **(H)** RTqPCR analysis of *NTHL1* isoforms. Isoform 1 exhibited the highest expression levels, while isoforms 2 and 3 were detected at lower levels. **(I)** gel electrophoresis of PCR using specific primers targeting each individual isoform. **(J)** gel electrophoresis PCR using primers able of amplifying all three isoforms. Isoform 1 exhibited the highest expression levels, while isoforms 2 and 3 were detected at lower levels. **(K)** Quantification of experiment depicted in J.

**Figure S2**

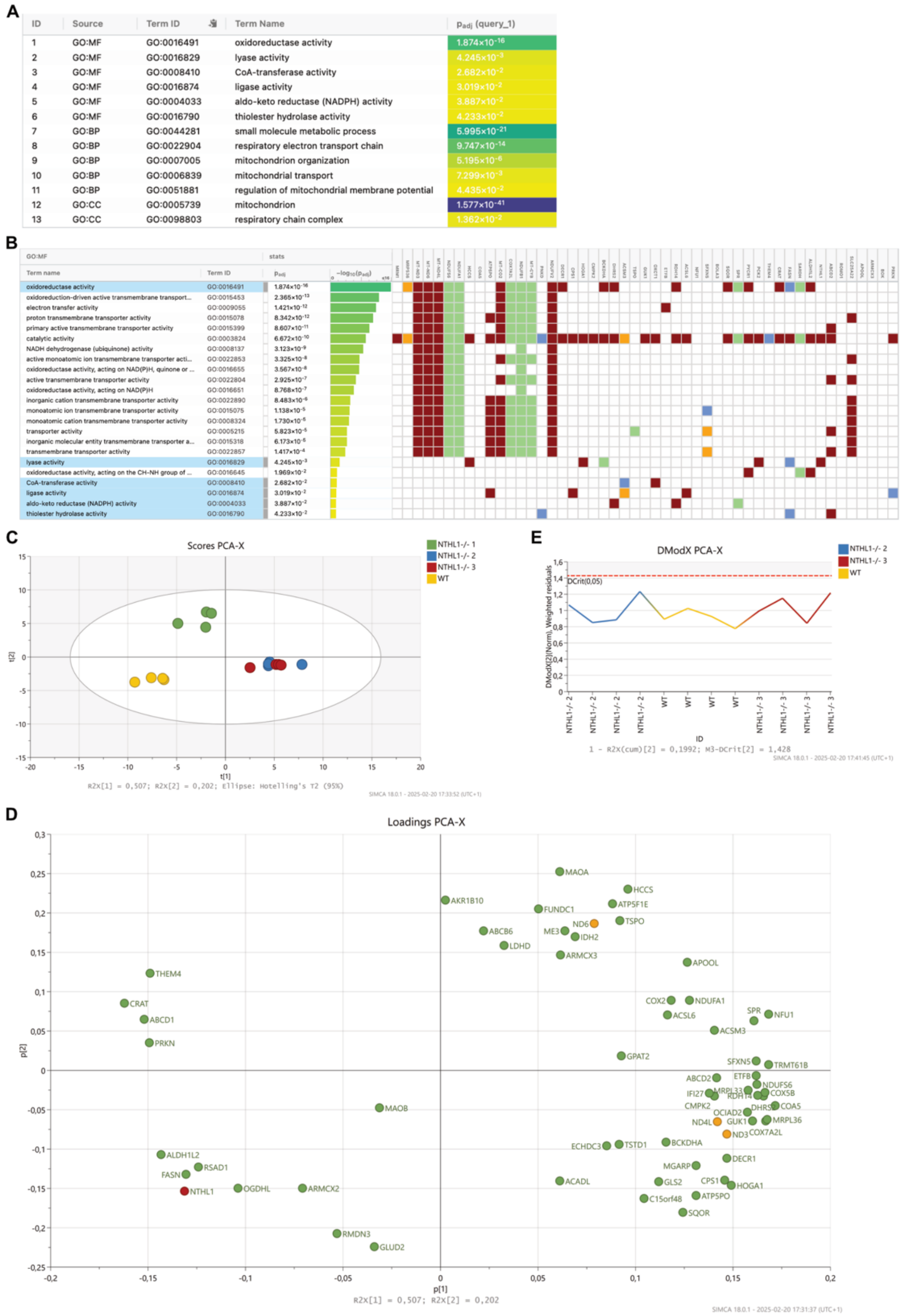


**Figure S2: GO Enrichment and OPLS-DA analysis of differentially expressed mitochondrial genes in *NTHL1-/-* and WT samples.** **(A)** GO Enrichment Analysis: top significantly enriched terms for molecular function (MF), biological process (BP), cellular component (CC) and their adjusted p-value (-$\log_{10}$ Padj). **(B)** GO Enrichment analysis of the top significantly enriched MF terms. The bar chart illustrates the gene set's primary contributions to key MF, with each bar representing a distinct GO category. Color intensity corresponds to statistical significance (adjusted p-value). **(C)** Score plot showing clear separation between *NTHL1-/-* 1 (green), 2 (blue), 3 (red) and WT (yellow) samples along the first predictive component (t[1]), indicating distinct expression profiles of mitochondrial genes between the two groups. **(D)** Loadings plot identifying the contributions of individual genes to the separation between KO and WT samples. **(E)** DModX line plot indicating no significant outliers in the dataset, with all samples falling below the critical threshold (DCrit = 0.05), confirming the robustness of the analysis.

### Figure S3

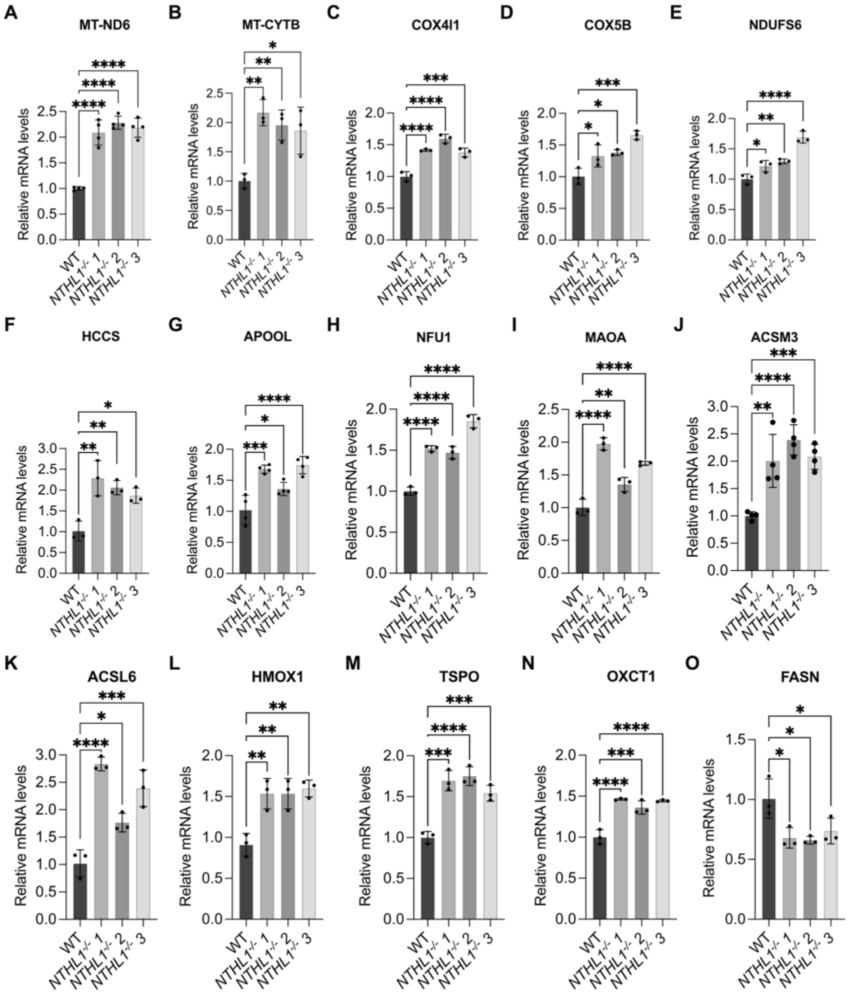


**Figure S3: RT-qPCR validation of RNA-Seq data.** RT-qPCR analysis confirms the differential expression of selected genes *MT-ND6* (A), *MT-CYTB* (B), *COX4l1* (C), *COX5B* (D), *NDUFS6* (E), *HCCS* (F), *APOOL* (G), *NFU* (H), *MAOA* (I), *ACSM3* (J), *ACSL6* (K), *HMOX1* (L), *TSPO* (M), *OXCT1* (N), *FASN* (O). Data are represented as fold change in $NTHL1^{-/-}$ cells relative to WT cells. Error bars represent mean ± SD ($n \geq 3$). *$p \leq 0.001$; **$p \leq 0.01$; ****$p \leq 0.0001$, one-way ANOVA.

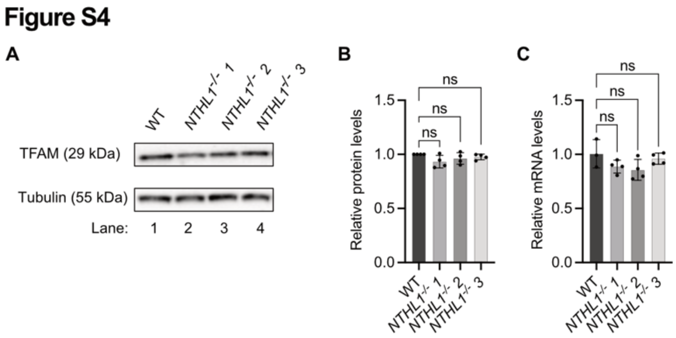


**Figure S4: TFAM expression is not affected in *NTHL1*$^{-/-}$ cell Lines. (A)** Immunoblot analysis of TFAM in extracts from HEK293 WT (lane 1) and 3 independent HEK293 *NTHL1*$^{-/-}$ clones (lanes 2-4). **(B)** Quantification of the data shown in panel A. **(C)** RT-qPCR Validation of RNA-seq Data: RT-qPCR analysis confirms no differential expression of selected TFAM gene. Data are represented as fold change in *NTHL1*$^{-/-}$ cells relative to WT cells. Error bars represent mean ± SD (n ≥ 3). one-way ANOVA. ns – not significant.

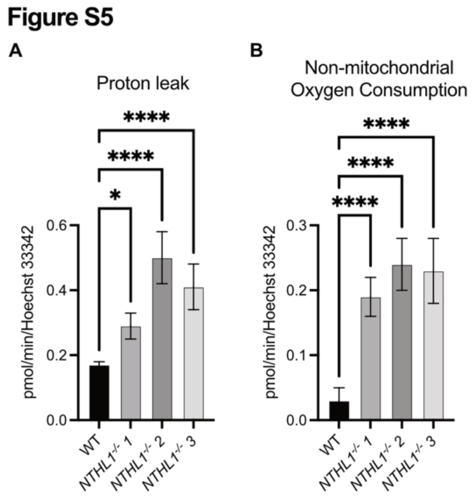


**Figure S5: Increased proton leak and non-mitochondrial oxygen consumption in NTHL1-deficient cell lines. (A)** Proton Leak: *NTHL1*-deficient cells show increased proton leak, indicating energy dissipation as heat instead of ATP production, suggesting altered mitochondrial efficiency. **(B)** Non-Mitochondrial Oxygen Consumption: These cells also exhibit increased non-mitochondrial oxygen consumption, reflecting heightened oxidative activity and potential ROS generation outside the mitochondria.

Figure S6

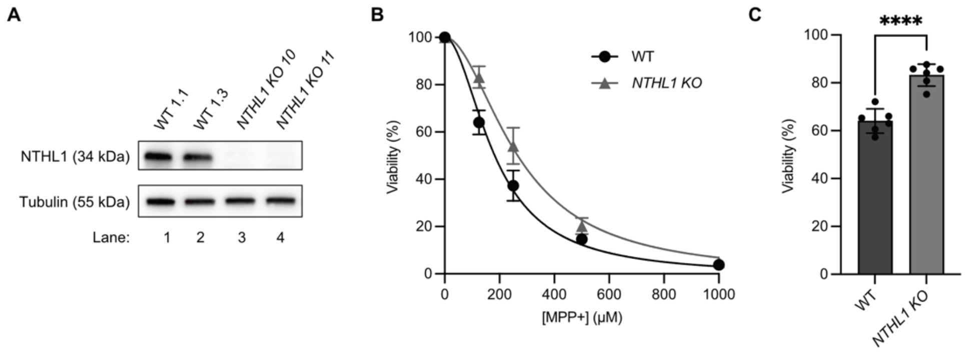


**Figure S6: *NTHL1*-deficient HAP1 cells display enhanced tolerance to 1-methyl-4 phenylpyridinium (MPP+). (A)** Immunoblot analysis of NTHL1 in extracts from HAP1 WT (lane 1, 2) and 2 independent HAP1 *NTHL1 KO* clones (lanes 3, 4). **(B)** Viability of HAP1 WT and *NTHL1 KO* cell lines after 72h treatment with indicated amounts of MPP+. **(C)** Viability of HAP1 WT and *NTHL1 KO* cell lines after 72h treatment with indicated 125 μM MPP+. Data are represented as fold change in *NTHL1*$^{-/-}$ cells relative to WT cells. Error bars represent mean ± SD (n ≥ 3). ****p ≤ 0.0001, t-test
